# Improved sensory representations as a result of temporal adaptation

**DOI:** 10.1101/2024.07.26.605075

**Authors:** Amber Marijn Brands, Zilan Oz, Nikolina Vukšić, Paulo Ortiz, Iris Isabelle Anna Groen

## Abstract

Human perception is robust under challenging conditions, for example when sensory inputs change over time. Temporal adaptation in the form of reduced responses to repeated external stimuli is ubiquitously observed in the brain, yet it remains unclear how repetition suppression aids recognition of novel inputs. To clarify this, we collected behavioural and electrocorticography (EEG) measurements while human participants categorized objects embedded in visual noise patterns after first viewing these patterns in isolation, inducing adaptation to the noise stimulus. We furthermore manipulated the availability of object information in the visual input by varying the contrast of the noise-embedded objects. Our results provide convergent behavioral, neural and computational evidence of a benefit of temporal adaptation on sensory representations. Adapting to a noise pattern resulted in overall faster object recognition and better recognition of objects as object contrast increased. These adaptation-induced behavioral improvements were accompanied by more pronounced contrast-dependent modulation of object-evoked EEG responses, and better decoding of object information from EEG activity. To identify potential neural computations mediating the benefits of temporal adaptation on object recognition, we equipped task-optimized deep convolutional neural networks (DCNNs) with different candidate mechanisms to adjust network activations over time. DCNNs with intrinsic adaptation mechanisms, such as additive suppression, best captured contrast-dependent human performance benefits, whilst also showing improved object decoding as a result of adaptation. Finally, adaptation effects in networks that use temporal divisive normalization, a biologically-plausible canonical neural computation, were most robust to spatial shifts, suggesting that temporal adaptation via divisive normalization aids stable representations of time-varying visual inputs. Overall, our results demonstrate how temporal adaptation improves sensory representations and identify candidate neural computations mediating these effects.

**Author summary:** Robust perception is essential for the human brain to detect, process, and act upon new sensory inputs. Temporal adaptation is believed to play a key role in robust sensory processing by allowing neurons to continuously adjust their responses to previous inputs in order to optimize the processing of future inputs. Here, we show that temporal adaptation aids visual object recognition by improving neural representations of object contrast and object category. By emulating temporal adaptation in deep convolutional neural network models with different computational mechanisms, we identify candidate neural computations mediating benefits of temporal adaptation on sensory processing.

## Introduction

In natural, real-world environments, humans often need to process sensory stimuli in challenging settings, for example when detecting an approaching car on a misty road (**Fig. 1A**). How does the human brain compute robust sensory representations in such suboptimal, dynamically varying viewing conditions? One neural phenomenon that may aid robust perception is temporal adaptation to previously perceived inputs. Numerous studies have shown that repeated sensory stimulation reduces neural responses, both in the visual modality (Grill-Spector et al. 2006; Henson 2003; Groen et al. 2022), and other senses (Whitmire and Stanley, 2016). Typically, the brain shows stronger response reductions for more similar inputs (e.g., Sawamura et al. 2006; Brands et al. 2024; **Fig. 1B**). This form of temporal adaptation, known as repetition suppression, can improve recognition of the repeated stimulus itself, a process known as priming (Desimone, 1996; Schacter and Buckner, 1998; Henson, 2003). However, adaptation is also thought to increase sensitivity to other, novel stimuli (the approaching car), by decreasing the saliency of recently seen stimuli (the misty road), so as to efficiently process changes in the environment (Barlow, 1993; Vogels, 2016; Clifford et al., 2007; Kohn, 2007). Benefits of adaptation on low-level vision (contrast sensitivity, orientation tuning and motion perception) have been extensively documented (e.g., Solomon and Kohn 2014), but its influence on higher-level neural representations, and how this aids perceptual performance, is less understood. Here, we examine how temporal adaptation facilitates sensory processing of objects by measuring human neural responses and recognition behavior with and without prior adaptation.

**Figure 1:**
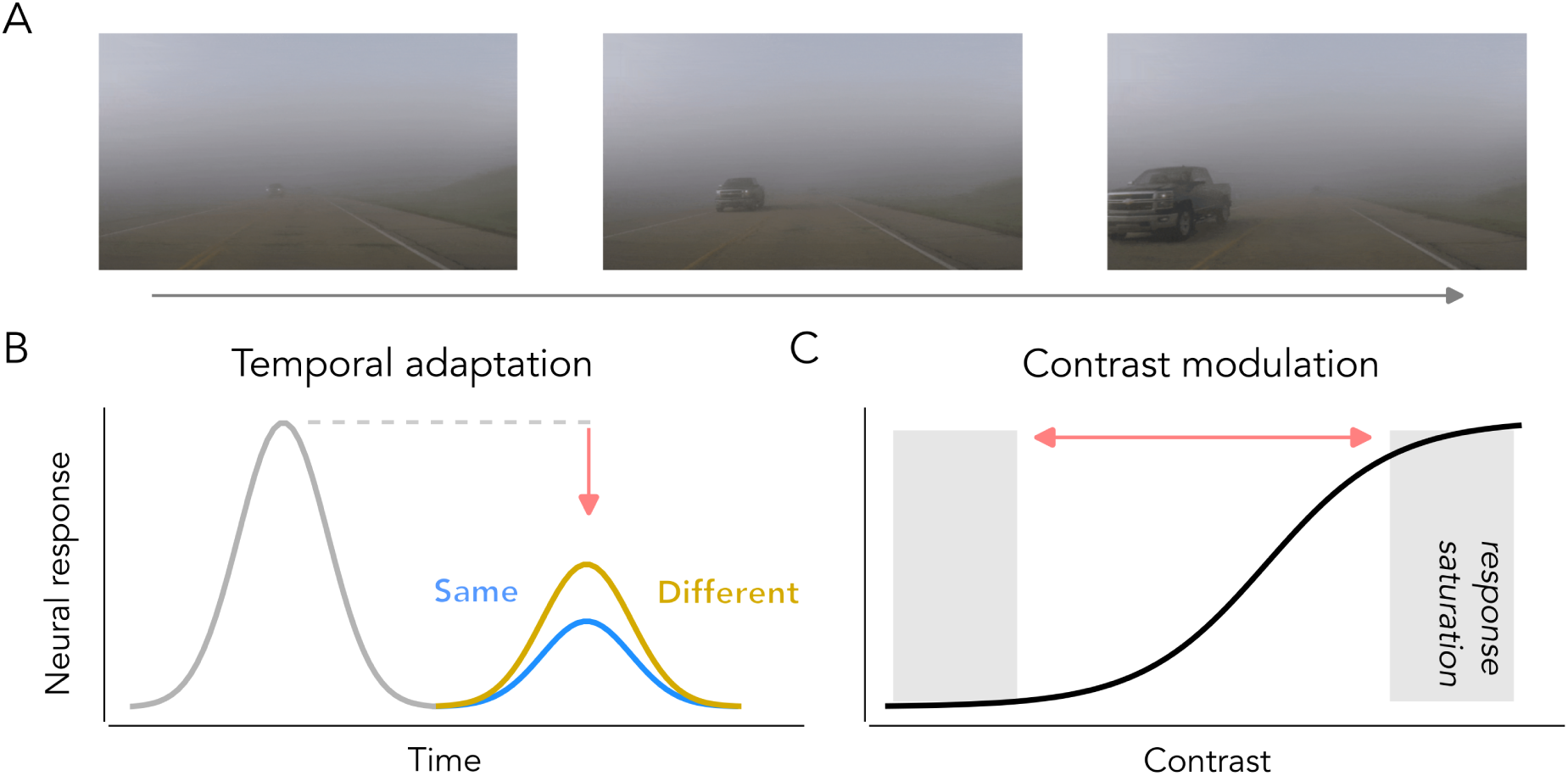
Neural response properties in visual cortex that affect perception under challenging viewing conditions. **A**: Humans are able to recognize temporally varying targets, for example an approaching car, in suboptimal viewing conditions, for example mist. **B**: Temporal adaptation refers to the reduction of neural responses when stimuli are repeated, with more pronounced reductions for similar compared to different repeated inputs. **C**: Contrast modulation is characterized by the contrast response function; neural responses are reduced for low input contrast.

Another outstanding question is what neural computations underlie temporal adaptation. Many neural response properties are thought to be governed by canonical computations, arising from intrinsic biophysical mechanisms operating within individual neurons (Whitmire and Stanley, 2016) or from recurrent interactions (del Mar Quiroga et al., 2016). A prior study by Vinken et al. (2020) investigated the computational mechanisms underlying benefits of adaptation on object recognition by endowing model neurons in a deep convolutional neural network (DCNN) with an additive adaptation state, and found that this simple intrinsic suppression mechanism was sufficient to mimic adaptation-induced object categorization improvements as observed in human participants. An alternative form of intrinsic adaptation, divisive normalization (Heeger, 1992, 1993), has been proposed as a canonical neural computation underlying a wide range of neural response properties (Carandini and Heeger, 2012), including repetition suppression in human intracranial recordings (Groen et al., 2022; Brands et al., 2024). Here, we investigate whether implementing divisive normalization in a DCNN offers an advantage over additive suppression in simulating improved object recognition after adaptation. Moreover, since prior work has shown that adding recurrent connections to a feedforward DCNN also capture carry-over effects in sequences of visual stimuli well (Tang et al., 2018; Sörensen et al., 2023), we also included a third adaptation mechanism based on lateral recurrence.

To elucidate how temporal adaptation to previously seen stimuli can aid novel object perception, human participants categorized objects embedded in repeated noise patterns, while electrocorticography (EEG) was measured. To manipulate task difficulty and better emulate naturalistic viewing conditions, we additionally varied the contrast of the target objects. Contrast reliably modulates visual response magnitudes, as characterized by the contrast response function (CRF) (Albrecht and Hamilton 1982; Albrecht et al. 2002; **Fig. 1C**). Although such modulations have been most thoroughly characterized in low-level visual areas, they also occur in higher-level areas, resulting in remarkably similar response reductions in intracranial responses as repeated, full-contrast stimuli (Groen et al., 2022). Here, we asked how these two types of response modulations interact and how they could potentially facilitate or compensate for one another. Based on the results by Vinken et al. (2020), we predicted that adapting to the noise pattern would reduce its saliency by suppressing responses to these stimuli, resulting in stronger responses to the hidden objects, thereby enhancing the quality of the object representations. However, since contrast reduction also suppresses neural responses, we expect temporal adaptation to benefit object representation in a graded fashion - more so for high than for lower object contrasts.

Our EEG analyses focused on characterizing the joint effects of noise adaptation and object contrast on object-evoked responses (ERPs), which are known to be sensitive to both stimulus repetition (Summerfield et al., 2011; Schweinberger and Neumann, 2016) and contrast (Vassilev et al., 1994; Schadow et al., 2007). Given that divisive normalization can account for both contrast- and repetition-induced response reductions (Groen et al., 2022), we anticipated that DCNNs with divisive normalization would more accurately capture potential interaction effects of repetition suppression and contrast on object recognition as compared to the additive model introduced by Vinken et al. (2020). We furthermore asked whether adaptation resulted in better decoding of object properties from the EEG signal, as well as convolutional neural network activations.

Consistent with joint effects of adaptation and contrast modulation, we find that temporal adaptation to the noise pattern leads to improved categorization performance for higher but not lower contrast objects. The behavioral improvement of adaptation on object recognition is accompanied by stronger contrast modulation in evoked responses and improved decoding of object category from EEG signals after adaptation. We find that DCNNs with intrinsic adaptation mechanisms more accurately capture human behavior and neural responses than lateral recurrence mechanisms. Moreover, DCNNs that employ intrinsic adaptation via divisive normalization are also more robust to spatial shifts of the adapter, suggesting that this mechanism contributes to temporal stability of adaptation. All together, these results suggest that robust object recognition arises from an interplay between temporal adaptation and contrast modulation and suggest that divisive normalization is not only an effective mechanism for capturing neural responses in sensory cortex, but also leads to perceptual robustness.

## Results

To examine how temporal adaptation improves representations of novel sensory inputs, we collected EEG data while participants categorized test images containing objects with various contrast levels embedded in noise patterns (**Fig. 2A**), after inducing distinct forms of adaptation. Specifically, we varied the adapter preceding the test image to be either the same of a different noise pattern (**Fig. 2B**), or a uniform control (blank; **Fig. 2C**). Below, we first reveal how prior adaptation to the noise pattern impacts object recognition performance, after which we show how it affects the representation of object class and contrast in event-related responses (ERPs) to the test images. Next, we compare several computational mechanisms that could underlie temporal adaptation using deep convolutional neural network (DCNN) modeling (**Fig. 2D**), and asses which models most accurately capture human object categorization performance and neural representations. Among the mechanisms tested are those inspired by previous work by Vinken et al. (2020), including an intrinsic suppression mechanism and a mechanism operating across feature maps through lateral recurrence (**Fig. 2E**). Additionally, we also examined whether an alternative, biologically-plausible form of temporal adaptation, namely divisive normalization, offers advantages over the other two mechanisms and better aligns with the human data (**Fig. 2F**). Lastly, we investigate the robustness of the DCNNs implemented with different adaptation mechanisms, particularly their ability to maintain performance when spatial shifts are applied to the input.

**Figure 2:**
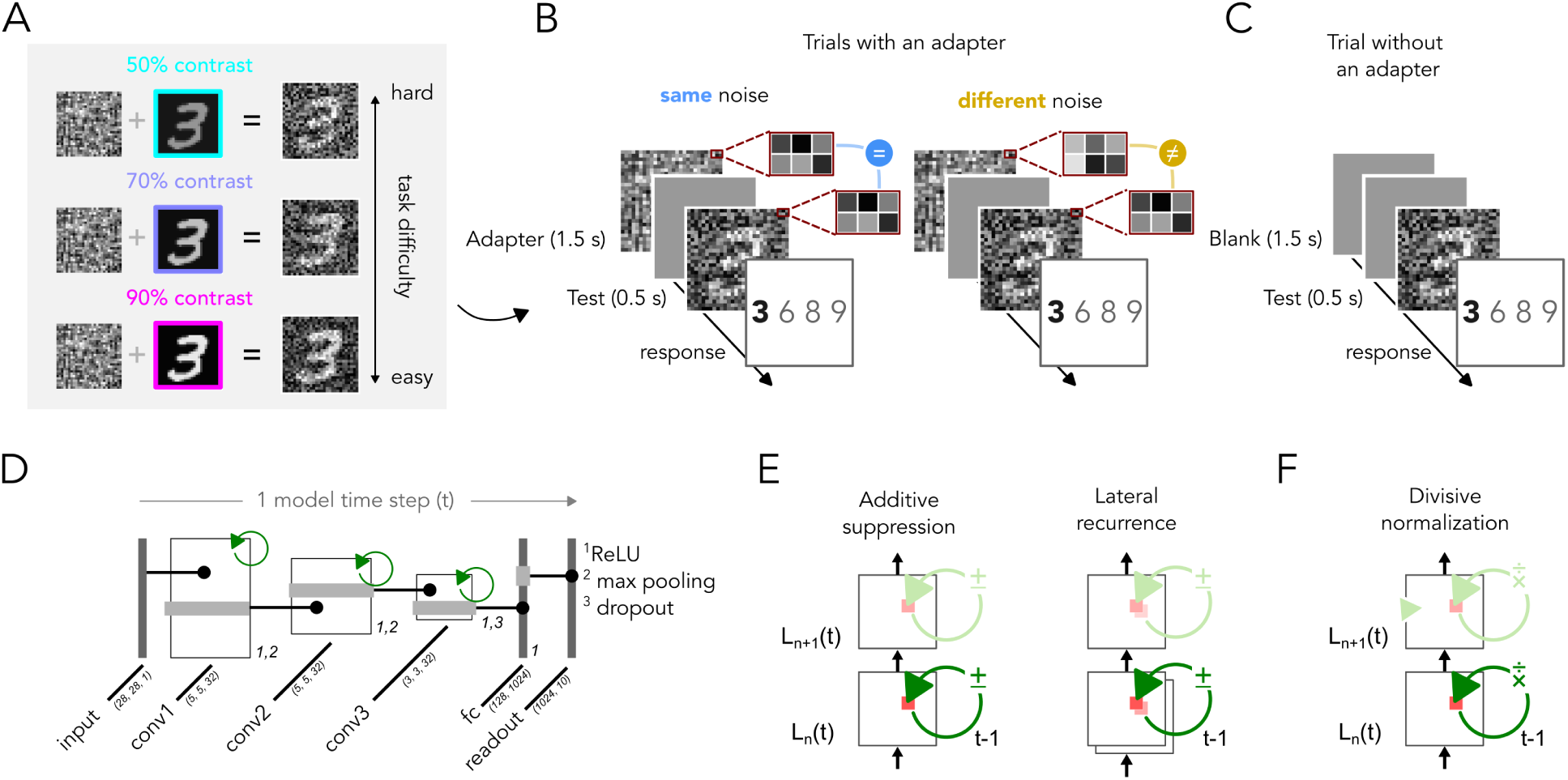
Experimental design and network modeling. **A**: Object recognition task with the test images consisting of objects (digits 3, 6, 8 and 9 from the MNIST dataset) embedded in a pixelised noise pattern. Contrast of the digit image was varied (50%, 60%, 70%, 80% and 90%). **B**: Adaptation trials, consisting of the presentation of same (left) or different (right) noise prior to the test image. **C**: Control trials, consisting of the presentation of the test image in isolation. **D**: DCNNs were trained with a feedforward backbone consisting of three convolutional filter layers, one fully connected layer and a readout layer. After each convolution, history-dependent adaptation was applied (depicted in green), feeding activations from the previous model time step, i.e. previous feedforward pass. Filter sizes are represented within parentheses. **E**: Temporal adaptation mechanisms as implemented by Vinken et al. (2020). *Left*, Additive suppression whereby each unit feeds its activations from the previous timestep. *Right*, Temporal adaptation mechanism which feeds unit activations across feature maps. **F**: Divisive normalization introduced by Heeger (1992, 1993) which feeds activations from the previous timestep using divisive rather than additive suppression, here implemented using a recursive multiplicative feedback signal (see Methods: DCNN modeling).

### Faster and more accurate object recognition after noise adaptation

First, we established that adding noise to the test images impaired object recognition accuracy: whereas subjects readily recognized clean objects (mean categorization performance = 98.15%, SD = 0.30%), performance decreased substantially for objects embedded in noise patterns (mean = 60.61%, SD = 9.39%), with stronger deterioration as object contrast decreased (**Fig. 3A**). We analyzed categorization accuracy using a linear mixed effects model (see Materials and Methods, *Statistical testing for behavioral data*) that tested for effects of adapter type (blank, same and different) and object contrast (50%, 60%, 70%, 80% and 90%). We found a main effect of adapter (*F* _(2_,_4)_ = 28.24, *p <* 0.001) and object contrast (*F* _(2_,_4)_ = 141.81, *p <* 0.001), as well as an interaction (*F* _(2_,_8)_ = 2.75, *p* = 0.006), indicating that adaptation differently affects performance across object contrasts. Adapting to noise had no effect for the two lowest contrast levels (50% and 60%), with similar performance across conditions, while for 70% object contrast, performance improved after adapting to both same and different noise, but not to the blank adapter (same, *p <* 0.001; different, *p <* 0.001). For the highest contrast levels (80% and 90%), adapting to the same noise resulted in higher performance compared to both blank (80%, *p <* 0.001; 90%, *p <* 0.001) and different noise adapters (80%, *p* = 0.04; 90%, *p* = 0.0019). These results confirm earlier findings showing a clear benefit of temporal adaptation on novel object recognition: the performance-degrading effect of the spatial noise patterns is diminished by the prior adaptation phase, with same-noise adaptation resulting in the most robust categorization improvement. Notably, adaptation did not improve recognition for the weakest object contrasts, indicating that human behavioral benefits of adaptation are somewhat contrast-dependent.

**Figure 3:**
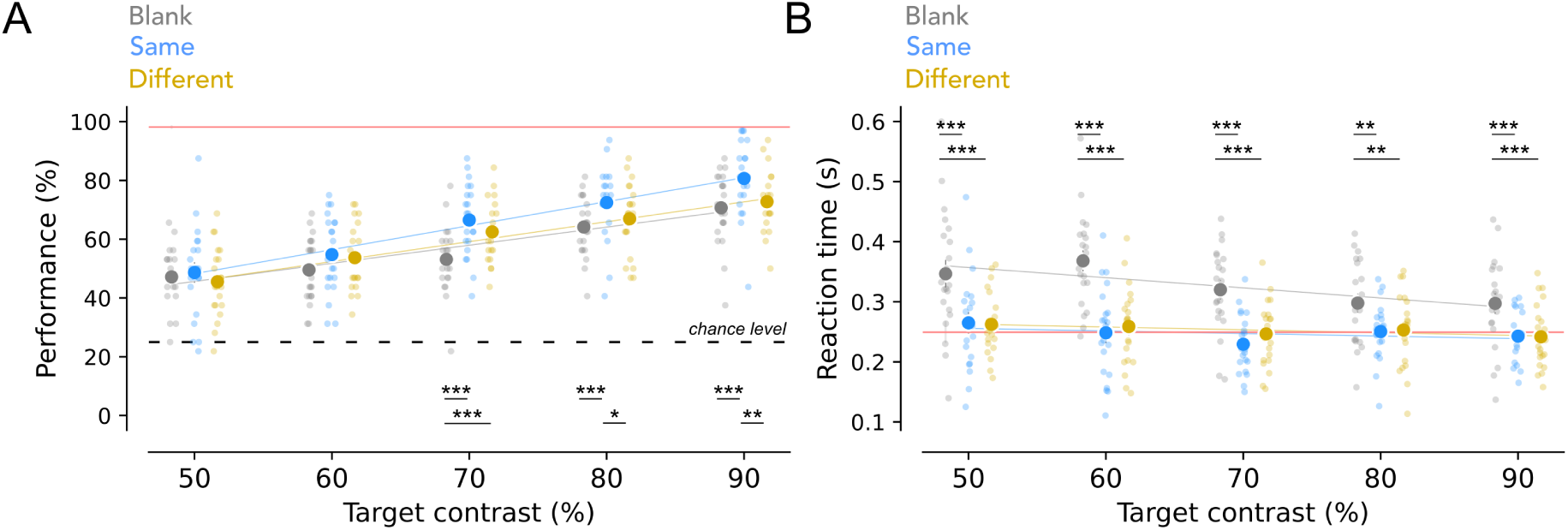
Effects of temporal adaptation on categorization performance and reaction times for noise-embedded objects. **A**: Accuracy across contrast levels for test images shown after a blank adapter (grey) or a same (blue) or different (yellow) noise adapter as the test image. The solid red line shows performance for objects without noise; the dotted black line shows chance level (25%). Each point depicts an individual subject. **B**: Similar as (A) but for reaction times. Adaptation to same noise most improves recognition performance for higher contrast levels, while adaption to both same and different noise results in faster reaction times overall. Linear mixed effect model, post-hoc Tukey test, ^∗^ p *<* 0.05, ^∗∗^ p *<* 0.01, ^∗∗∗^ p *<* 0.001. This figure can be reproduced by mkFigure3.py.

We also examined how temporal adaptation affected reaction times (RTs) to the test images. Similar to categorization accuracy, we found a main effect of adapter (F_(2_,_4)_ = 105.09, *p <* 0.001) and contrast level (*F* _(2_,_4)_ = 7.42, *p <* 0.001) as well as an interaction effect (*F* _(2_,_8)_ = 2.81, *p* = 0.005). However, the pattern of RT differences between conditions was different from that observed for accuracy; response times for test images presented with a noise adapter (either the same or different noise) were significantly shorter compared to blank adapters for all object contrasts (pairwise *p <* 0.001), with largest RT effects for the lowest contrasts (**Fig. 3B**). In fact, having a noise adapter resulted in RTs similar to those of clean object images, suggesting that temporal adaption relieved the detrimental impact of the added noise on response speed. Together, our behavioral results reveal clear benefits of temporal adaptation on object categorization performance, with improved recognition of noise-embedded objects (given sufficient object contrast) after first adapting to the same preceding noise, accompanied by overall faster responses after adaptation.

### Temporal adaptation affects early and late responses in occipito-parietal electrodes

We next examined how visually-evoked ERPs were modulated by temporal adaptation. All test images, regardless of the presented adapter, elicited deflections in occipital and parietal electrodes both early (100-150 ms, **Fig. 4A**, *top*) and later in the response (∼ 300 ms, **Fig. 4A**, *top*). Direct comparison shows that ERP differences due to adaptation were most pronounced in occipito-parietal electrodes, evident in a clear difference in ERP amplitude between the test images with noise vs. blank adapters (**Fig. 4B**). Notably, responses after same-noise adaptation showed the largest deflection difference compared to the blank adapter. To quantify these response differences, we first determined which time points showed significant effects of repetition suppression, by testing when the average ERP to the noise-adapted test images differed from those to test images following a uniform control adapter (see Materials and Methods, *Identification of time windows of interest*). We then computed ERP response magnitudes separately for same- and different-noise adapters in those time windows. For occipital electrodes we found two significant time windows with suppression effects (45-111 ms and 201-271 ms), but these contained no significant difference between same and different noise adapters (**Fig. 4C**). For parietal electrodes, we found three time windows, whereby similar to occipital electrodes, the first two (29-80 ms and 158-228 ms) did not differ across adapter types, but during the third time window (341-431 ms), same-noise adapters resulted in larger ERP deflections than different noise adapters (dependent-samples T-test, *t* _(20)_ = −3.54, *p* = 0.002, **Fig. 4D**). These results show that temporal adaptation to a noise pattern affects evoked activity to test images in both occipital and parietal electrodes, with adapter-specific effects emerging later in time at parietal electrodes only. There, same-noise adaptation results in stronger ERP deflections than different-noise adaptation, possibly reflecting facilitated neural processing of the object due to repetition suppression of the noise adapter.

**Figure 4:**
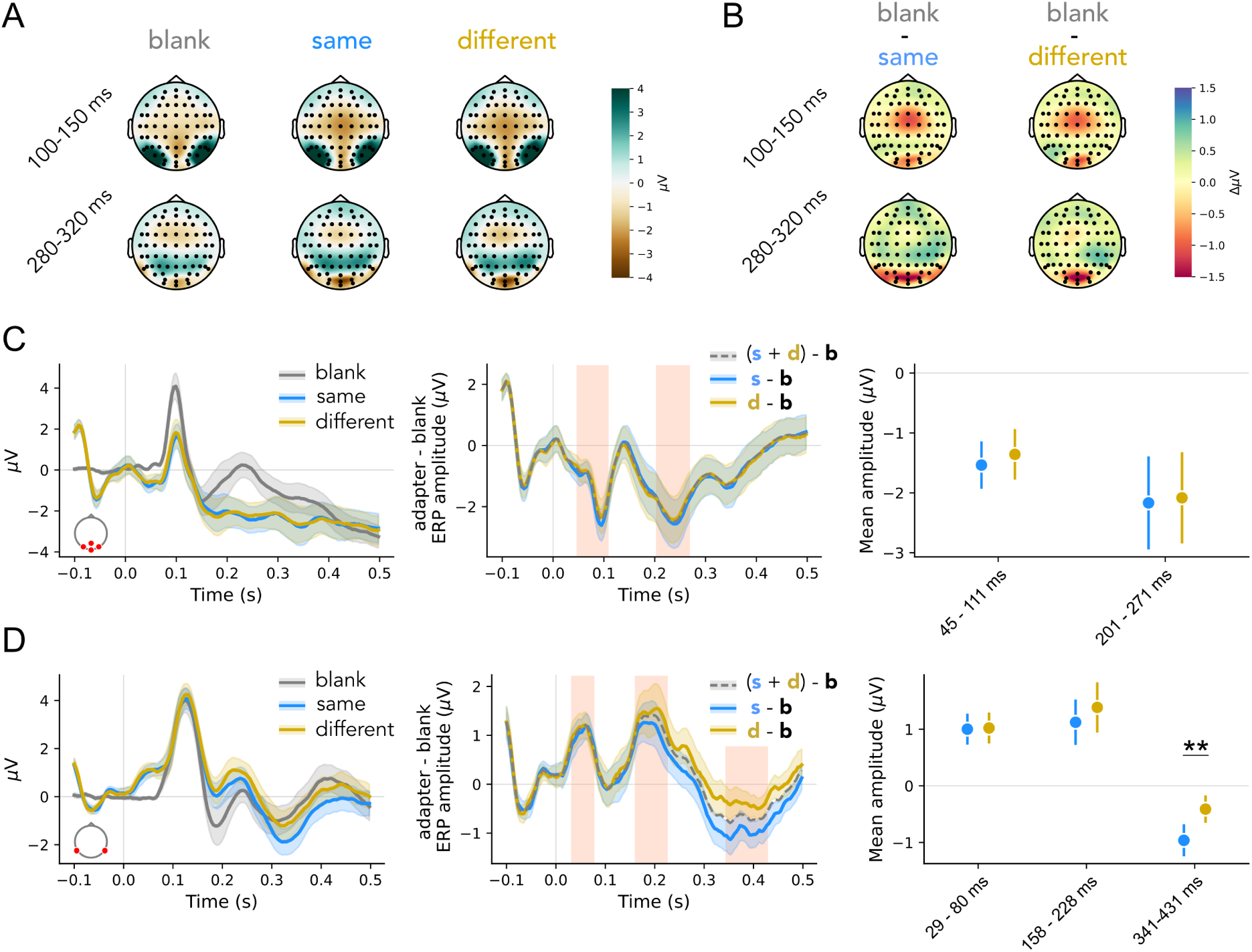
Temporal adaptation results in enhanced object-evoked activity at parieto-occipital electrodes. **A**: Topomaps showing ERP amplitudes for test images presented without adapter (blank, left) and test images presented after adapting to same (middle) or different noise (right). **B**: Difference in evoked potentials to test images with and without adapters (left, same noise versus blank; right, different noise versus blank). **C**: Left, average ERP per condition for electrodes Iz, Oz, O1 and O2. Middle, differences in ERPs with and without adaptation. Time windows for test images with adaptation for which same- (s) and different-noise (d) conditions are significantly different from blank (b) conditions are depicted by shaded red areas. Right, average ERP amplitudes for same vs. different noise adaptation within the identified time windows. Shaded regions and errorbars depict SEM across subjects. **D**: Same as C, but for electrodes P9 and P10. Adaptation-specific differences arise in parietal electrodes later in the response, with stronger deflections when adapting to same as opposed to different noise. T-test (two-sided), ^∗∗^ p *<* 0.01. This figure can be reproduced by mkFigure456.py.

### (Occipito)-parietal electrodes exhibit contrast-dependent response modulation

We have shown that evoked responses are modulated by preceding inputs and that adaptation-specific differences occur later in the response. In addition to inducing temporal adaptation, we also varied the contrast of the noise-embedded objects. Consistent with the fact that the test images differed only marginally in their overall contrast level (which was dominated by the contrast of the noise pattern; see Materials and Methods, *Stimuli*), we found no object contrast-dependent differences in early ERP responses (*<* 200 ms, **Fig. 5A**). Interestingly, however, later responses (*>* 200 ms) in both occipito-parietal and parietal, but not occipital channels varied systematically with object contrast (**Fig. 5B-D**). To test for an effect of object contrast, we performed a linear fit on the average ERP amplitude in the late time window (see Materials and Methods, *Identification of time windows of interest*) and found a significant non-zero slope at parietal (one-sample T-test, slope of linear fit against 0, *t* _(20)_ = −8.93, *p <* 0.001, **Fig. 5B**) and occipito-parietal (*t* _(20)_ = 6.24, *p <* 0.001, **Fig. 5C**), but not occipital electrodes (*t* _(20)_ = −0.51, *p* = 0.62, **Fig. 5D**). These results show that response amplitudes of parietal electrodes, in addition to being modulated by temporal adaptation, were also modulated by the contrast of the noise-embedded object.

**Figure 5:**
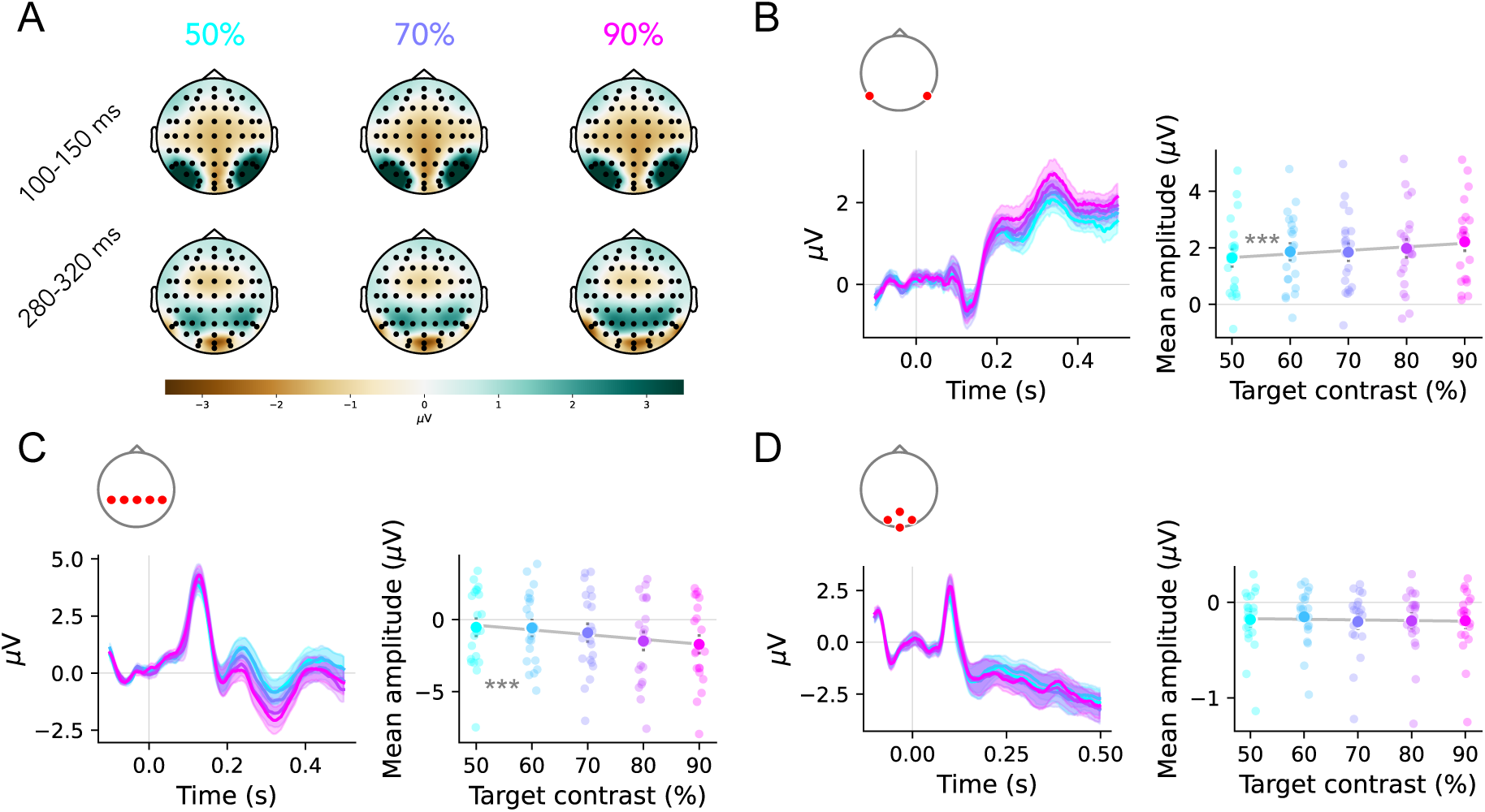
Responses in (occipito-)parietal, but not occipital, electrodes are modulated by the contrast level of noise-embedded objects. **A**: Topomaps representing ERPs during presentation of the test image for three object contrast levels. **B**: Left, ERPs shown separately per contrast level for P9 and P10. Right, response magnitude computed by taking the mean amplitude for the P300 component. **C-D**: Same as B for occipito-parietal (C) electrodes, including Pz, P1, P2, P3 and P4, and occipital electrodes (D), including Iz, Oz, O1 and O2. The P300 component of (occipito-)parietal electrodes is modulated by object contrast. 1-sample T-Test (coefficient of linear curve against 0), ^∗∗∗^ p *<* 0.001. This figure can be reproduced by mkFigure456.py.

### Increased contrast-dependence after temporal adaptation in parietal electrodes

So far, we found contrast-dependent benefits of temporal adaptation on object categorization performance, as well as independent effects of noise adaptation and contrast on ERPs. To test whether temporal adaptation in fact facilitated neural object processing, we next searched for evidence of enhanced object contrast processing as a consequence of adaptation. Since ERP effects of temporal adaptation and object contrast co-occurred on parietal electrodes (P9 and P10), we specifically focus on these electrodes to test for the interaction, by now separating responses to test images with different object contrast levels by the type of adapter. This more focused comparison indicated that parietal electrodes indeed exhibited more pronounced contrast-dependent modulation after same-noise adapters, compared to blank or different-noise adapters (**Fig. 6A**). A linear fit on response amplitudes in late time windows across object contrast levels shows that adapting to the same noise results in significant contrast-dependent modulation (**Fig. 6B**), with increasing negative amplitudes for increasing contrast levels (one-sample T-test, slope of linear fit against 0, *t* _(20)_ = 3.32, *p* = 0.003). Although ERP responses to test images in the other conditions showed similar trends, the linear fit was not significant (different-noise trials, *t* _(20)_ = 0.57, *p* = 0.57); blank trials, *t* _(20)_ = 0.62, *p* = 0.54). Moreover, these adapter-specific effects on contrast modulation were not observed in the occipital and occipito-parietal electrode groups. To conclude, our EEG data reveal that temporal adaptation induces specific modulations of EEG responses to noise-embedded objects, resulting in more pronounced object contrast-dependent responses in occipito-parietal electrodes beyond 200 ms of visual processing.

**Figure 6:**
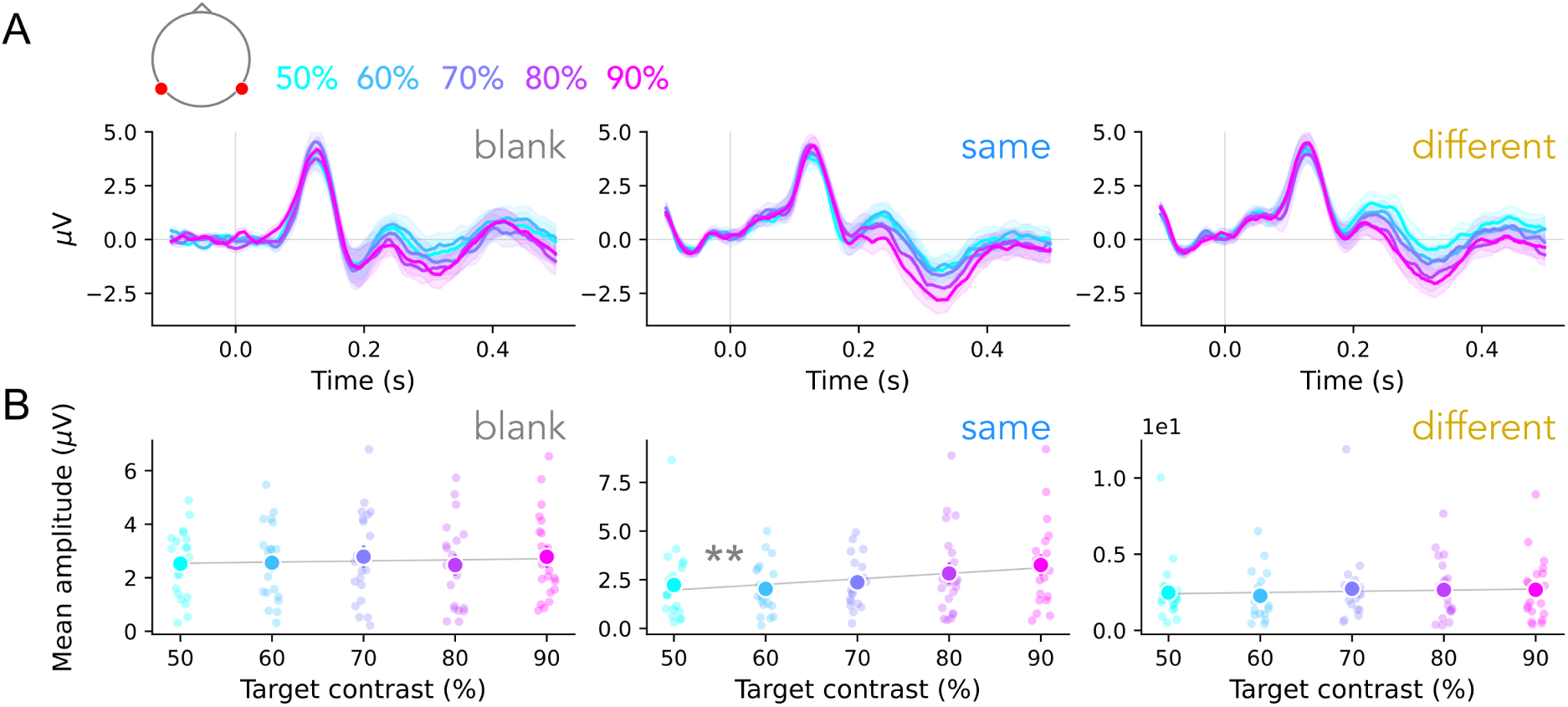
More pronounced object contrast modulation at parietal electrodes after same-noise adaptation. **A**: ERPs for P9 and P10 shown separately for trials without adaptation (left) and adaptation with same (middle) or different (right) noise compared to the test image. The shaded regions depict the SEM across subjects. **B**: Response magnitude was computed by taking the mean amplitude for the P300 component for the similar experimental conditions as in panel (A). ERP signals exhibit significant object contrast-dependent modulation, evident as increasingly negative response deflections for increasingly higher contrast levels, after adapting to same noise only. 1-sample T-Test (coefficient of linear curve against 0), ^∗∗^ p *<* 0.01. This figure can be reproduced by mkFigure456.py.

### Temporal adaptation leads to increased object decoding from ERP signals

The more pronounced neural modulations by object contrast after same-noise adaptation suggests that repetition suppression of the noise pattern resulted in better neural encoding of the noise-embedded object. While this increased contrast-dependence may indicate enhanced processing of the embedded objects, it does not directly demonstrate that same-noise adaptation benefits object recognition performance by improving the object discriminability. To test this, we assessed how well we could decode object class from ERP responses to the test images (see Materials and Methods, *ERP Decoding Analysis*). If temporal adaptation improves behavioral categorization by enhancing the neural representation of the object, we expect more successful decoding after same-noise adapters, compared to blank or different-noise adapters.

The results show that inducing temporal adaptation to the noise pattern indeed results in better object decoding from EEG responses. We first verified that object representations were detectable in the evoked activity by performing the decoding analysis on ‘clean’ objects presented in full contrast without embedded noise, which resulted in strong above-chance decoding accuracy (one-sample T-test, *t*_(20)_ = 8.48, *p <* 0.001; **Fig. 7A**). Unsurprisingly, embedding the objects in noise strongly reduced the quality of object representations, since for test images preceded by blank adapters, decoding accuracy dropped to chance level. Remarkably, however, these object-representations were partly restored after adaptation to noise, resulting in significant above-chance decoding accuracy after both same-noise (one-sample T-test against chance level, *t*_(20)_ = 3.36, *p* = 0.003) and different-noise adapters (one-sample T-test, *t*_(20)_ = 2.79, *p* = 0.01), mirroring the improved behavioral object recognition performance as a result of adaptation.

**Figure 7:**
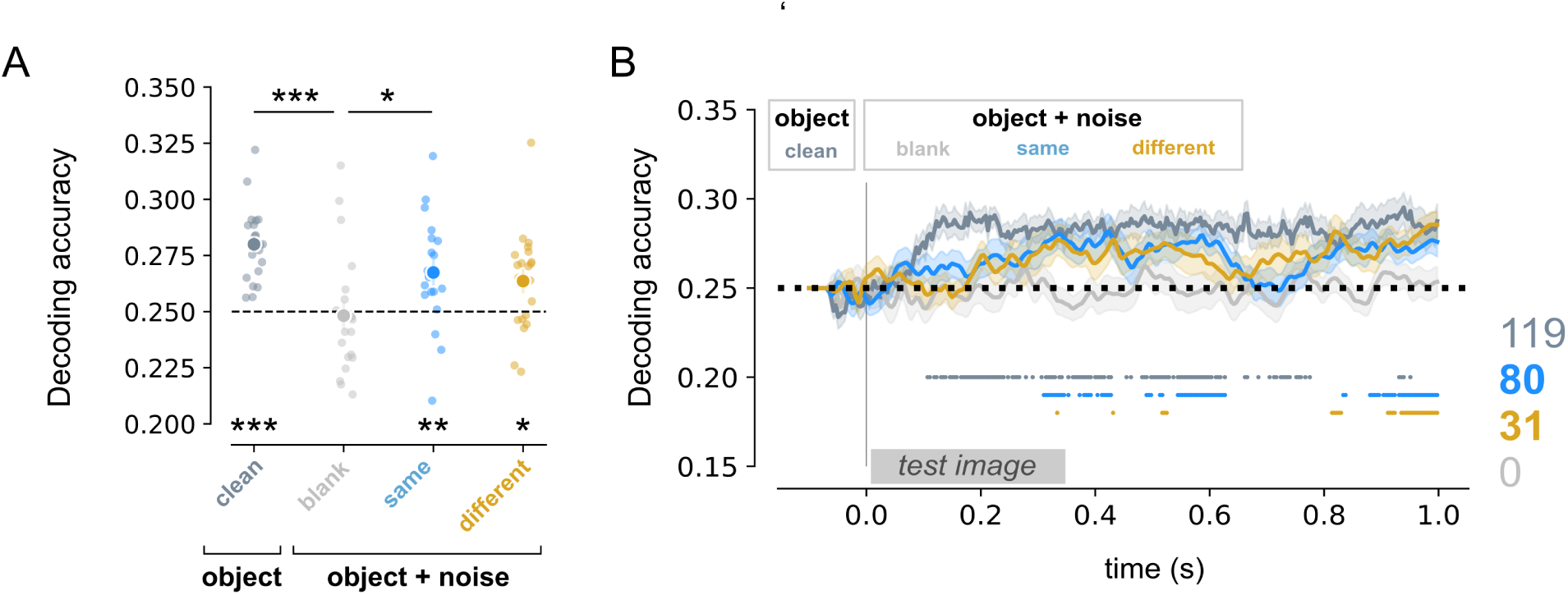
Temporal adaptation improves decoding of objects from EEG responses. **A**: Average decoding accuracy for predicting the presented object class based on evoked potentials for test images without (dark gray) and with embedded noise, including blank (light grey), same- (blue) and different-noise (orange) trials. Decoding accuracy’s are averaged for [0, 1]s time window after stimulus onset. The lower asteriks denote significant differences from chance level (0.25, one-sided T-test). The upper asteriks denote significant differences across trial types (one-way ANOVA, post-hoc Tukey test). ^∗^ p *<* 0.05, ^∗∗^ p *<* 0.01, ^∗∗∗^ p *<* 0.001. **B**: Decoding accuracy for the different trial types across time points, with the number of time points for which decoding accuracy was significantly different from chance level (i.e. 25%) noted on the right. Shaded regions depict the SEM across subjects. This figure can be reproduced by mkFigure7_SFig2.ipynb.

Statistical comparison between decoding accuracies for the different adaptation conditions indicated a main effect of adapter type (one-way ANOVA, *F* _(3)_ = 7.02, *p <* 0.001), with a significant difference in decoding performance between clean and blank trials (*p <* 0.001) and between same-noise and blank trials (*p* = 0.036), but not between different-noise and blank trials (*p* = 0.13), suggesting an adapter-specific effect on object-representations in the evoked activity. Indeed, decoding accuracy was significantly above chance for substantial more time-points during same-noise trials compared to different-noise trials (**Fig. 7B**), which aligns with the most robust perceptual benefit of same-noise adaptation observed in behavior.

We also assessed whether EEG object decoding accuracy was affected by object contrast. While we did not find a main effect of contrast on average decoding accuracy (**Supp. Fig. 2A**, one-way ANOVA, *F* _(4)_ = 0.34, *p* = 0.85), we did observe above chance decoding accuracy for substantially more time-points for higher than for lower contrast levels (50% = 10, 60% = 26, 70% = 64, 80% = 43, 90% = 76 significant time-points), consistent with the observed improved categorization accuracy in behavior (**Supp. Fig. 2B**). Overall, these results demonstrate that temporal adaptation enhances neural decoding of object information under challenging viewing conditions, suggesting that the behavioral benefit of temporal adaptation on object recognition is mediated by improvements in the underlying neural representation.

### DCNNs with a single-unit adaptation mechanism align better with human behavior and neural responses

Our results so far show that temporal adaptation to a noise pattern improves participants’ ability to recognize objects embedded in that same noise pattern, in particular for higher object contrasts (**Fig. 8A**), and that these behavioral improvements are accompanied by more distinct neural representations of object contrast and category (**Fig. 6-7**). To discern potential computational mechanisms that could mediate these perceptual and neural effects, we endowed DCNNs with three different adaptation mechanisms (**Fig. 2D-F**; see Materials and Methods, *DCNNs: temporal adaptation mechanisms*). Two of these, additive suppression and divisive normalization, operate on the level of the single model unit, whereby responsiveness to network inputs is reduced proportional to previous activations of only the unit itself. The other adaptation mechanism uses lateral recurrence, operating across feature maps from previous timesteps. By training and testing the DCNNs on the same task as our human participants (see Materials and Methods, *DCNN modeling: Neural network training*), we examined which mechanism best emulated the observed object recognition benefits of temporal adaptation in behavior and EEG. Following an earlier implementation (Vinken et al., 2020), our first batch of networks were trained on short sequences (*t* = 3), with the adapter and test image presented for one model timestep each. We also included a DCNN without any temporal adaptation as a baseline.

**Figure 8:**
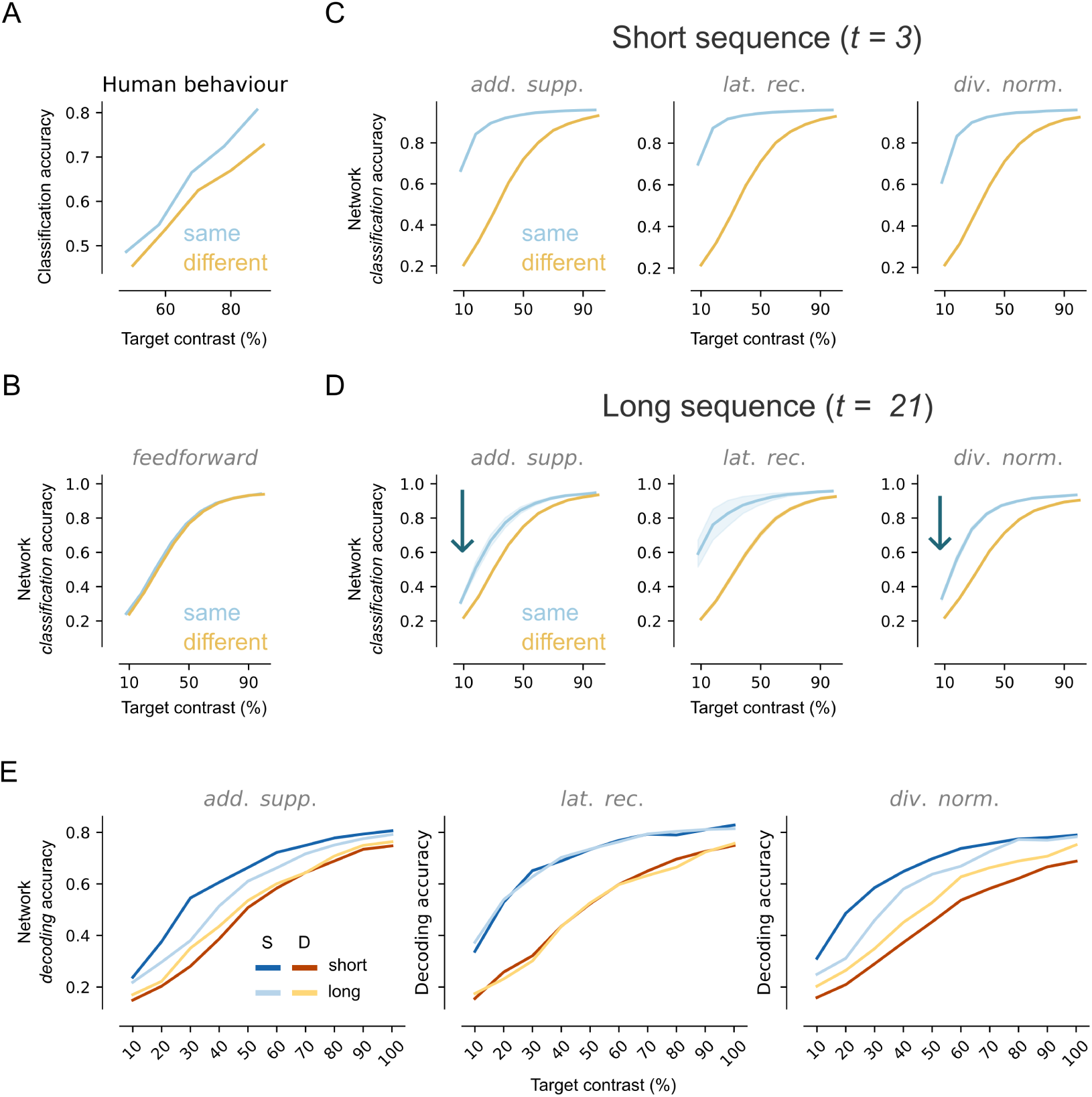
DCNNs with extended intrinsic adaptation show human-like benefits on performance and neural object representations. (*previous page*) **A**: Human object recognition performance for same- (blue) and different- (yellow) noise trials (same data as in Fig. 3A). **B**: Network classification accuracy for the test set after same- and different-noise adapters for DCNNs without a temporal adaptation mechanism. **C**: Network classification accuracy for DCNNs with one of three temporal adaptation mechanisms (from left to right): additive suppression, lateral recurrence and divisive normalization. Input sequences consisted of three images: an adapter (A), a blank (B) and a test (T) image (*ABT*). **D**: Same as panel C for networks trained on input sequences of 21 images (*AAAAAAAAAAAAAAABTTTTT*). Networks with instrinsic adaptation mechanisms better approximate the human behavior, showing increasing benefit of temporal adaptation for higher object contrasts. **E**: Decoding accuracy for the first convolutational layer for the test set for same (blue) and different (yellow) noise adapters for DCNNs with temporal adaptation. Decoding performance for later layers is shown in **Supp. Fig. 8**. Adapting to the same noise leads to better object decoding of for all adaptation mechanisms. Panels A-D can be reproduced by mkFigure8ABCD.py and panels E can be reproduced by mkFigure8E_SFig8.py.

As expected, the baseline model did not benefit from the presentation of the same noise prior to the test image, as this network processes each image independently (**Fig. 8B**). In contrast, all networks with temporal adaptation mechanisms show a benefit of adaptation, evident from higher DCNN classification performance for same compared to different noise trials (**Fig. 8C**), in line with human behavior (**Fig. 8A**) and previous modeling work (Vinken et al., 2020). However, all models failed to accurately predict the interaction between adapter type and contrast exhibited by humans, who showed stronger performance benefits for same-noise adaptation for high but not low object contrasts. In fact, the models showed the opposite pattern, namely a larger benefit of adaptation for lower rather than higher contrast levels. We reasoned that this mismatch could be due to the short training sequence we used: in our human experiment, the adaptation phase lasted for several seconds, allowing temporal accumulation of adaptation (Akyürek, 2025), which may be necessary for the emergence of contrast-dependent modulation of neural responses (Ohzawa et al., 1985). In contrast, the DCNN models could only process the input images for a single time step each, prohibiting prolonged adaptation. Therefore, we trained a second batch of models on increasingly longer input sequences, allowing the models to fully exploit their adaptation mechanisms and to better approximate the extended adaptation period in the human experiment.

Optimization on longer input sequences revealed that DCNNs with temporal adaptation indeed more closely mimic human behavior, evident by the fact that the benefit of adaptation decreased for lower, but not higher object contrasts (**Fig. 8D**). Interestingly, the interaction between contrast and temporal adaptation observed in human behavioral performance was better matched by models with intrinsic adaptation mechanisms (additive suppression and divisive normalization) than models with lateral recurrence, which continued to show quite a large improvement for same-noise adaptation even for low-contrast objects. Moreover, unit activations in DCNNs with lateral recurrence did not reduce, but rather increased during the adaptation period, whereas DCNNs with intrinsic adaptation mechanisms showed clear reductions; and while all three models exhibited repetition suppression and contrast modulations to test images, divisive normalization models showed most stable temporal dynamics across initializations (**Supp. Fig. 3-7**).

Finally, we also tested whether DCNNs with adaptation mechanisms similarly exhibit more separable object representations in their model activations, following adaptation. To test this, we performed an analogous object class decoding analysis as for the EEG responses on the model unit activations, separately for each DCNN model layer (see Materials and Methods, *DCNN modeling: DCNN decoding analysis*). The results show that already in the first model layer (**Fig. 8E**; results for other layers are provided in **Supp. Fig. 8**), all three types of models indeed demonstrated improved object decoding for same compared to different noise adapters. Moreover, DCNNs with intrinsinc adaptation, but not lateral recurrence, again better mirrored the human data, in the form of more improved object decoding for higher but not lower contrasts, after prolonged adaptation.

Overall, our comparison between DCNNs with human behavioral and EEG response patterns suggests that intrinsic adaptation mechanisms best account for the observed perceptual benefits of temporal adaptation on human object categorization behavior, provided the model has sufficient temporal input for the adaptation mechanism to take effect.

### Adaptation in DCNNs with divisive normalization is more robust to spatial shifts

Our computational modeling analysis found that DCNNs with intrinsic adaptation mechanisms yielded highest similarity with human behavior, showing neurally plausible response reductions during the adaptation period. While more similar to human behavior and neural decoding across object contrasts, these networks were however objectively *worse* than networks with lateral recurrence in terms of overall performance. Could temporally extended adaptation via intrinsic mechanisms potentially benefit other aspects of perception, such as robustness or representational stability? Compared to humans, classic DCNNs show poorer robustness and generalisation to distorted images (Geirhos et al., 2018b), with small image perturbations resulting in large deviations in model behavior, a property that is famously exploited in adversarial network attacks (Szegedy et al., 2013). To investigate whether temporal adaptation can enhance network robustness, we ran a new version of our adaptation experiment on the DCNN models whereby we introduced minor spatial perturbations on the noise patterns. Specifically, we used the same noise pattern for both adapter and test image, but now spatially shifted the pixel values of the latter, such that the resulting test image has a slight offset. We then assessed to what extent the benefits of adaptation on object recognition persisted across these small perturbations, and to what extent this depended on the length of the adaptation period.

Reflecting DCNNs sensitivity to pixel-level changes, shifting the noise pattern during test indeed resulted in reduced accuracy (compared to no shift) for all the temporal adaptation mechanisms (**Fig. 9A**), indicating that spatial perturbations reduce the benefit of adaptation for object recognition in these models. However, robustness of adaptation increased as sequence length increased, in particular so for DCNNs with divisive normalization. In fact, we find a main effect (one-way ANOVA, *F* = 40.99, *p <* 0.001) with DCNNs employing divisive normalization showing significant increases in performance with more prolonger adaptation, unlike DCNNs with additive suppression (one-way ANOVA, *F* = 1.24, *p* = 0.33) and lateral recurrence (one-way ANOVA, *F* = 0.46, *p* = 0.77). Moreover, DCNNs with divisive normalization show the least performance reduction over individual pixel shifts with prolonged adaptation (**Fig. 9B**), suggesting this mechanism maintains the adapter-induced suppression most robustly, resulting in better performance at test time. Overall, these results demonstrate that DCNNs with divisive normalization not only accurately capture benefits of temporal adaptation on neural object representations and object recognition behavior as observed in human behavior, but also show more spatially robust adaptation, supporting its role as a biologically plausible adaptation mechanism supporting stable sensory representations.

**Figure 9:**
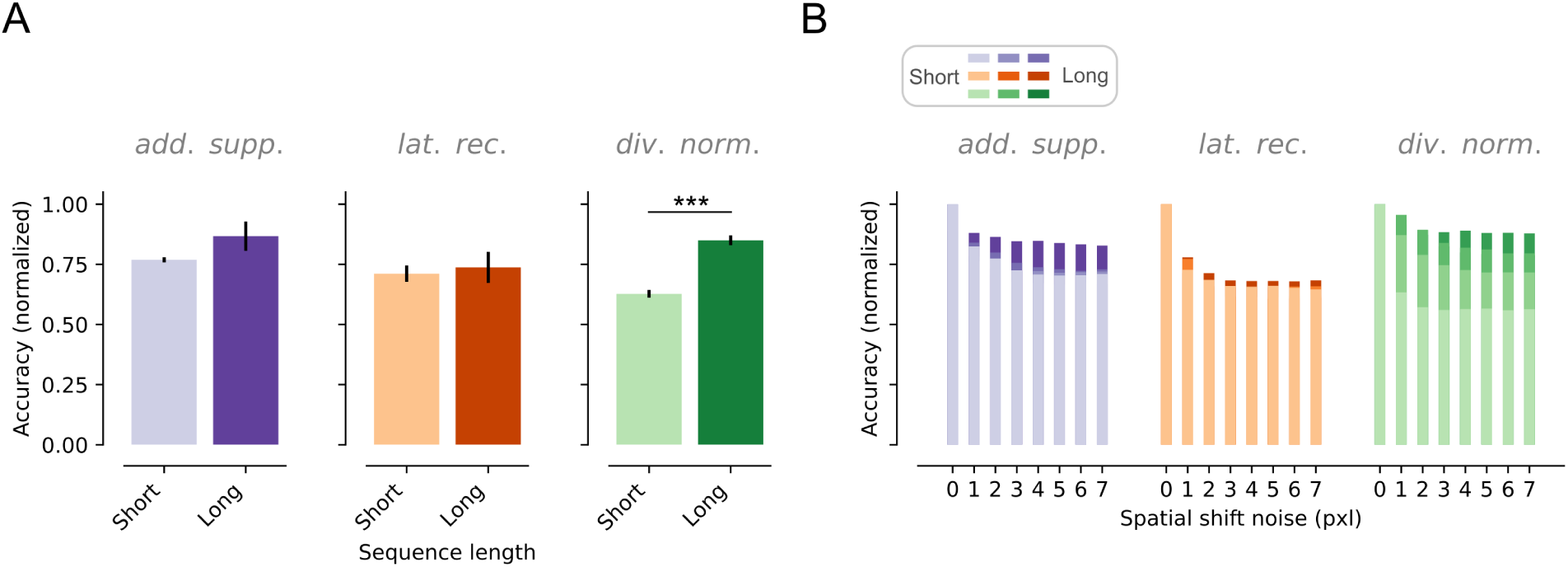
DCNNs with divisive normalization show higher robustness against spatial shifts of input. **A**: Average accuracy across spatial shifts for DCNNs with temporal adaptation trained on short (n = 3, ABT) or long (n = 21, *AAAAAAAAAAAAAAABTTTTT*) input sequences. The accuracy is normalized with respect to no shift. DCNNs with divisive normalization (*DN*) are more robust against spatially shifting noise during test time as the sequence length increases compared to DCNNs with additive suppression (*AS*) and lateral recurrence (*LR*). Error bars show the SEM. Independent T-test, ^∗∗∗^ p *<* 0.001. **B**: Effect of spatially shifting the noise during the test image on network performance. Depicted is the drop in accuracy compared to 0 pixel shift for the three temporal adaptation mechanisms optimized on the different sequence lengths. This figure can be reproduced by mkFigure9.py.

## Discussion

Our aim was to examine how temporal adaptation aids object recognition in human visual cortex. Confirming prior work, we find that inducing temporal adaptation to a visual noise pattern leads to improved subsequent recognition of objects embedded in that same noise pattern. Here, we show that this adaptation-induced behavioral improvement is accompanied by more pronounced modulation of high-level EEG responses by object contrast, as well as improved decoding of the object class from EEG. Together, these results show that temporal adaptation improves neural representations of noise-embedded objects by allowing the brain to reduce the representation of the surrounding noise, resulting in more robust representation of the object.

We also examined which computations could potentially explain the benefit of temporal adaptation by implementing several candidate mechanisms in DCNNs. We demonstrate that DCNNs with intrinsic mechanisms operating in individual model units (i.e. lacking lateral interactions) most accurately capture the behavioral and neural effects observed in humans. Moreover, DCNNs that adapt via intrinsic divisive normalization show better robustness to spatial shifts of the adapter. Overall, these findings suggest that improved object recognition due to temporal adaptation may result from relatively simple, intrinsic mechanisms, while pointing to temporal divisive normalization in particular as a promising biologically plausible mechanism to increase temporal stability of object representations in convolutional neural networks.

### The role of temporal adaptation on object recognition behavior

Previous work has suggested that temporal adaptation could serve to decrease salience of recently seen stimuli (Solomon and Kohn, 2014; Vogels, 2016). We demonstrated this principle by testing human observers and DCNNs on a task with temporally repeated, but task-irrelevant noise patterns, whereby reducing the salience of recently seen features resulted in perceptual changes evident by increased categorization performance. In addition to temporal adaptation, we studied the effects of contrast-dependent modulation of neural responses, which are characterized by a contrast response function (Albrecht and Hamilton, 1982). The benefit of temporal adaptation was only apparent for objects with sufficient contrast, suggesting that the saliency reduction of the noise patterns could not fully compensate for lowered contrast of the object. Interestingly, we found faster reaction times when participants were presented with an adapter regardless of object contrast, and regardless of whether the adapter noise was the same or different to the test image. This dissociation between human performance and reaction time suggests that different neural mechanisms may underlie improvements in categorization accuracy versus processing speed. The faster reaction times could potentially be attributed to a phenomenon known as contrast gain, referring to a shift of the dynamic range of the CRF as a result of pre-exposure to our high-contrast adapters (Wilson and Humanski, 1993; Dao et al., 2006), which has been previously shown to indeed affect reaction times (Cao and Pokorny, 2010). Since the contrast of the same- and different noise adapters were identical, however, contrast gain cannot explain the specific benefits of same-noise adaptation on categorization accuracy - this effect can rather be attributed to repetition suppression. Overall, these findings emphasize the importance of studying neural response properties simultaneously, as is the case in more naturalistic settings, to better understand how their combined effects influence different facets of perception, including performance and processing speed.

### Signatures of repetition suppression and contrast gain in evoked neural responses

Our findings are consistent with prior EEG results showing effects of temporal adaptation (Garrido et al., 2009; Engell and McCarthy, 2014) and contrast-dependent modulation (Schadow et al., 2007; Groen et al., 2013; Xi et al., 2020) on evoked responses. Our EEG measurements show clear evidence of repetition suppression, in the form of reduced responses to the repeated noise pattern early in time, but increased responses later in time after adapting to same compared to different noise, suggestive of enhanced processing of the object stimulus due to repetition suppression. At these late time-points, we also found specific effects of the object contrast, with larger deflections as the object contrast level increases. More interestingly, we also observed a joint effect of temporal adaptation and object contrast, with stronger contrast-dependent modulation when adapting to same compared to different or no noise. This finding suggests that the shift of the CRF due to pre-exposure to the high contrast adapter, in combination with the response reduction induced by adapting to the same noise, together result in a larger gain of the neural response to the object, resulting in better categorization performance. An earlier study by Chaumon and Busch (2014) showed that a reduction of the gain in EEG responses results in reduced accuracy during a detection task, suggesting a link between the input-output-relationship and visual performance. It is important to note that our observations are correlational: future research should investigate in more detail the possibly causal relationship between the gain in neural response magnitude and categorization performance. Nonetheless, we show that the joint effects of temporal adaptation and contrast gain on categorization performance are reflected in the evoked responses, revealing a link between neural processing and perception during object recognition.

### Effective implementation of temporal adaptation in DCNNs

Consistent with the results by Vinken et al. (2020), we found that DCNNs with temporal adaptation operating on the level of the single unit best capture behavioral adaptation in our noise adaptation task. In addition, we show these DCNNs capture the interaction between temporal adaptation and contrast gain on categorization performance, but only when we increased the temporal integration window with which the networks could deploy their adaptation mechanism. Importantly, achieving the better match with human behavior necessitated lower network accuracy for low-contrast objects, thus lowering overall object recognition performance. This could mirror a trade-off in sensory adaptation: while adaptation can help filter out redundant information, it may also reduce sensitivity to weak but behaviorally relevant signals when the system has already attenuated similar input over time. The DCNNs’ ability to mimic this balance suggests that single unit-level mechanisms are sufficient to approximate the trade-off effects of adaptation underlying human perception. This computational framework provides a promising approach to investigate how adaptation shapes object representations in the brain in time-varying environments.

In addition to mechanisms operating on the single-unit level, we implemented one form of recurrence operating across channels. While these networks capture the behavioral benefit of adaptation for higher contrast levels, they were not affected by sequence length and failed to capture the interaction between temporal adaptation and contrast modulation. These results may inform us about which perceptual effects could arise from visual processing within individual neurons and which arise due to recurrent interactions. Given the abundance of recurrence in the visual system (Felleman and Van Essen, 1991; Sporns and Zwi, 2004; Markov et al., 2014) it is most likely that temporal adaptation is implemented by a combination of intrinsic cell properties and recurrent connections. Future work could include other tasks or extend the current model combining both intrinsic adaptation properties and lateral and/or top-down recurrence (Spoerer et al., 2017; Thorat et al., 2021; Lindsay et al., 2022). Nonetheless, we demonstrate how the role of nonlinear neural response properties in perceptual adaptation and object recognition can be elucidated using image-computable DCNNs, an approach that aligns with the neuroconnectionism program (Doerig et al., 2023) which uses artificial neural networks to model behavior and neural information processing.

### Decoding object representations from EEG responses and DCNN activations

We show that high-contrast objects presented without noise can be decoded from EEG responses, which is consistent with prior studies (Contini et al., 2017; Grootswagers et al., 2019). More interestingly, we observe an effect of temporal adaptation, where the drop in decoding accuracy as a result of embedding objects in noise can be recovered by adapting to the same or a different noise pattern. While it has been previously shown that stimulus adaptation can improve decoding of objects from neural responses in monkey IT (Kaliukhovich et al., 2013), to our knowledge facilitatory effects of prior adaptation on object decoding from EEG signals have not been previously reported. We also show that adaptation to same noise results in higher decoding accuracies of the object class for more timepoints compared to adaptation to different noise, suggesting that the perceptual benefit of temporal adaptation may be mediated by improved discriminability of object-related neural representations. DCNNs with temporal adaptation modulated by object contrast show a similar pattern as the neural data, with higher decoding accuracies from unit activations for same-compared to different-noise trials. This convergence in decoding behaviour between humans and DCNNs further supports the idea that temporal adaptation enhances the separability of object representations under challenging conditions. Overall, we reveal a correspondence between informative features representing object identity in evoked responses and DCNN activations, linking neural activations patterns to perceptual experience.

### Temporal divisive normalization enhances temporal stability of adaptation

We offer a potential advantage for divisive normalization over other implementations of temporal adaptation, evident by the fact that DCNNs endowed with this adaptation mechanism are more robust against spatially shifting inputs compared to networks with additive suppression and lateral recurrence. One explanation for this difference in robustness could be the multiplicative manner in which divisive normalization feeds back previous inputs as opposed to the additive computations used by the other two mechanisms. Multiplicative feedback effectively scales the current response based on prior activity, adjusting the gain of neural signals, whereas additive feedback simply shifts the response by adding or subtracting a fixed amount. By scaling responses relative to prior activity, this computation may better preserve the relative strength of feature representations, thereby making object identity more resilient to spatial distortions. Previous work has similarly observed benefits of multiplicative interactions, showing improved object recognition of images consisting of composite images of multiple categories (Konkle and Alvarez, 2024). It has also been suggested that multiplicative effects play a role in shaping the CRF during attention-related processes (Liu et al., 2021) and are a useful inductive bias for conditional computations (Jayakumar et al., 2020). Overall, characterizing the unique role of different adaptation mechanisms in how information flow is conditioned is a promising avenue and valuable addition to recent endeavors taken to explore network robustness (e.g. Geirhos et al. 2018a).

### Limitations and future work

First, during the object recognition task, object contrast was varied within a fixed range across subjects. Previous work however, has shown large heterogeneity in contrast sensitivity across individuals (Baker and Graf, 2009). Consequently, it is likely the lowest and highest contrast conditions yielded different neural response variations depending on subject-specific CRFs. Future studies could determine more fine-grained contrast ranges on the individual subject level, such that effects of temporal adaptation in dynamical as well as saturating contrast ranges can be investigated. Second, in the current study neural responses were collected with EEG. It is however unclear how CRFs of individual neurons translate to population responses. While in our EEG data analysis we quantify contrast-dependent modulation using a linear fit (**Fig. 6B**), previous studies have shown nonlinear IO relationships between visual inputs and neural responses (Chan et al., 2022). To further investigate how contrast-dependent modulation affects neural responses and how this leads to perceptual experience in humans, other brain measurements could be deployed such as intracranial EEG, which can be measured more locally and whose high-frequency content is more closely related to spiking activity (Miller et al., 2009; Ray and Maunsell, 2011) and behavior (Miller et al., 2014). Lastly, there are several possible extensions to the current modeling framework, including testing on additional tasks (e.g. object occlusion, multiple-object recognition) and computational mechanisms (e.g. power law adaptation,; Sörensen et al. 2023) to further study the role of various canonical computations and resulting response dynamics in robust object recognition.

## Materials and Methods

### Participants

Twenty-four subjects participated in this study (age mean ± SD, 21 ± 2 years, 2 males), which was approved by the Ethical Committee of the Psychology Department at the University of Amsterdam. All participants gave written informed consent before participation and were rewarded with study credits or financial compensation (10 euro/h). Two participants were excluded from analysis because of incomplete recordings, and one participant was excluded because of technical issues that arose during the recording session, yielding a total of 21 included participants.

### Stimuli

We adapted the object categorization task used in Vinken et al. (2020), who used hand-drawn doodles hidden in a temporally repeated noise pattern. Here, we extended this paradigm by manipulating object contrast, whilst replacing the hand-drawn doodles with digits. Participants were instructed to categorize exemplars of four digit classes (3, 6, 8 and 9) from the MNIST dataset (LeCun et al. 2010) presented with varying object contrast levels (50%, 60%, 70%, 80% and 90%) (**Fig. 2A**). The digit was embedded in noise, which consisted of adding a pixelized grey-scale noise pattern which was drawn from a uniform distribution on interval [0, 1) following a pink distribution (such that the power spectral density was inversely proportional to the frequency of the signal). The image consisting of the noise-embedded digit will from here on be referred to as the *test* image. Varying the contrast of the digit resulted in easy (high contrast) and difficult (low contrast) task conditions, while only marginally affecting the overall contrast of the test image, which was mainly determined by the noise pattern (Root Mean Square test image, mean 50% = 0.45, SD = 0.018; mean 60% = 0.46, SD = 0.018; mean 70% = 0.47, SD = 0.019; mean 80% = 0.48, SD = 0.024; mean 90% = 0.49, SD = 0.026). Depending on the trial type (see below), participants were also presented with *adapter* images consisting of noise patterns without embedded digits, as well as uniform grayscale control adapters.

### Experimental procedure

Participants completed one EEG recording session lasting approximately 90 minutes, consisting of two blocks with a total of 880 trials. Participants were instructed to fixate on a red cross in the center of a grey screen throughout each trial. Categorization responses were acquired by keyboard presses and inter-trial intervals were sampled from a uniform distribution between 1.25 and 1.75 seconds. After every 50 trials, a break was inserted during which participants were encouraged to take a short rest.

During the first block, participants were presented with three different trial types. Two trial types consisted of the presentation of an adapter (1.5 s), followed by a short grayscale interval and then the test image (0.5 s), with the adapter containing the same noise as that of the subsequent test image or a randomly generated different noise pattern (**Fig. 2B**). We refer to these trials where same or different noise was presented prior to the test image as *same-noise* and *different-noise* trials, respectively. The grayscale interval between the adapter and test was fixed (134 ms) and for each trial, a different object exemplar was selected and a unique noise pattern was created, such that all stimuli were trial-unique and there were no repetitions across trials. The third trial type served as a control - referred to as the *blank* trial - which consisted of a blank image followed by the presentation of the test image (**Fig. 2C**). For the *same-noise*, *different-noise* and *blank* trials eight exemplars per digit were presented for each contrast condition, resulting in a total of 480 trials (160 per adapter type, i.e. four targets × five contrast levels × eight exemplars).

In order to conduct a decoding analysis (see below) a second block of trials was presented at the end of the session during which participants categorized the target digits (100% contrast) without any added noise. Here, 100 exemplars were presented for each target digit class, resulting in a total of 400 trials.

### EEG acquisition and preprocessing

EEG recordings were made with a Biosemi 64-channel Active TwoEEG system (Biosemi Instrumentation; www.biosemi.com), using an extended 10–20 layout modified with two additional occipital electrodes (I1 and I2, while removing electrodes F5 and F6). Eye movements were monitored with electro-oculograms (EOGs). The EEG signal was digitized at 1024 Hz sampling rate. Recording was followed by offline referencing to external electrodes placed on the mastoids. Preprocessing for the purpose of computing event-related responses (ERPs) was done in Python, MNE (Gramfort et al., 2013), following standard pipelines in our lab (e.g., (Groen et al. 2012, see preprocessing createEpochs.ipynb): a high-pass filter at 0.01 Hz (6 dB/octave); a low-pass filter at 30 Hz (6 dB/octave) (Butterworth bandpass zero-phase, two-pass forward and reverse, non-causal filter; filter order 16, effective, after forward-backward; cutoffs at 0.01, 30.00 Hz: −6.02, −6.02 dB); a notch filter (zero-phase) at 50 Hz; epoch segmentation in −100 to 500 ms from stimulus onset of the test image; baseline correction between −100 and 0 ms; ocular correction using the EOG electrodes (Gratton et al., 1983); conversion to Current Source Density responses (Perrin et al., 1987).

After computing epochs for each of the 880 trials per subject, we removed epochs containing artifacts for individual electrodes and subjects (preprocessing epochSelection.py) as follows: We first computed the maximum response in the [-0.1, 0.5] time-window (voltage time courses were first full-wave rectified), after which the standard deviation (SD) of these maximum values over all trials was computed. Trials were excluded from analysis if the maximum response was *>* 2 SD. Based on the selection methods described above, on average 6.7% (min: 4.4%, max: 8.2%) of the epochs were excluded.

### Data analysis

#### Statistical testing for behavioral data

Categorization accuracy and reaction times were computed separately for each subject and trial-type, and subsequently fitted with linear mixed effects models (LMMs) using the *lme4* package implemented in R (Bates and DebRoy, 2004) (statistics lmer.R). We choose an LLM approach because humans are known to exhibit individual differences in contrast sensitivity (Baker and Graf, 2009), resulting in substantial variation in categorization performance across subjects. The model comprised two fixed effects: adapter type (blank, same or different) and contrast level (50%, 60%, 70%, 80% and 90%), and one random effect (subject). The estimated coefficients for each of the fixed effects were evaluated with ANOVAs, and the resulting p-values were corrected for multiple comparisons using Tukey method for comparing three estimates (i.e. the adapter types).

#### Averaging procedure for ERP analysis

In total, we obtained 880 single ERPs resulting from presentation of the test image, including 480 obtained from the noise-embedded object trials and 400 obtained from clean object trials. For the ERP analysis responses were averaged over trials as follows: For estimating the effect of adapter type, trials were averaged over exemplars, digit classes and contrast levels, yielding 160 ERPs per adaptation condition. For estimating the effect of contrast level, trials were averaged over exemplars, digit classes and adapter types, yielding 96 ERPs per contrast condition. For estimating the interaction effect between adapter type and contrast, trials were averaged over exemplars and digit classes yielding 32 ERPs per condition.

#### Identification of electrodes of interest

We focus our analyses on electrodes recording from early visual areas, including Oz, Iz, O1 and O2, and from higher-order visual areas, including P9 and P10, based on prior literature showing object-related processing/repetition suppression effects at these electrodes (Gruber and Müller, 2002; Schweinberger and Neumann, 2016; Johnsdorf et al., 2023). Moreover, for examining the contrast-dependent modulation, we also inspected parietal-occipital electrodes, including Pz, P1, P2, P3, and P4, which have been implicated contrast-related mechanisms (Di Russo et al., 2001). For the decoding analysis (see Modeling section below), single ERPs of all electrodes were used, motivated by the fact that this required no *a priori* assumptions regarding the spatial distribution of object-related representations.

#### Identification of time windows of interest

To study the effect of adapter types, we first determined which time windows were affected by preceding inputs. We achieved this by subtracting the ERPs of trials with blank adapters from the trials with same- or different-noise adapters. Subsequently, we determined the timepoints for which the noise-blank difference was significantly different from 0 using a one-sample t-test. To investigate the effect of contrast, we selected time windows implicated in early and late visual processing. The early time window included the P100, with an onset of 100-150 ms after stimulus onset, which has been linked to a first wave of image-selective neural activity facilitating core object recognition (Thorpe et al., 1996; DiCarlo and Maunsell, 2000). The late time window was centered on the P300 component (280-320 ms), which is known to reflect the engagement of higher-order cognitive functions, such as decision-making behavior during visual processing (Nazari et al., 2010; Philiastides and Sajda, 2006).

#### ERP decoding analysis

To examine how adaptation and object contrast jointly affect representations of object classes in noise in EEG responses, we ran a series of decoding analyses, which consisted of training a linear classifier to predict the presented digit (i.e. 3, 6, 8 or 9) based on the spatially-distributed ERP responses. We performed a subject-wise decoding analysis separately for the ERPs obtained by presenting either clean or noise-embedded test images (mkFigure8AB.ipynb). We used the Python sklearn function `sklearn`.`svm`.`SVC()`, with default parameters, which is able to handle binary as well as multi-class classification. To reduce computation time, ERP signals were first downsampled to 128 Hz. To increase the strength of the signal compared to the background noise a sliding window of 62.5 ms was used. For the trials presenting the noise-embedded test images (480 in total), a k-fold cross-validation was applied with 15 folds during which each ERP was used for inference once, with 448 images used for training and 32 images used for testing per fold. Here, the images used for testing within each fold belonged to one contrast level and one adapter type (e.g. same-noise adapter for an object contrast of 50%). For clean test images, the classifier was trained on 360 trials and tested on 40 trials. Similarly to the trials presenting the noise-embedded objects, a k-fold cross-validation was performed with 10 folds such that each trial was used once during inference.

### DCNN modeling

DCNN models were trained on the same object recognition task as the human participants, using stimulus sequencies containing both same and different noise images preceding the test image. We incorporated temporal dynamics into the networks by feeding the activity from previous timesteps, with one timestep, *t*, defined as one feedforward pass through the network, during which a single image is processed.

### Stimuli

In the original paradigm by (Vinken et al., 2020), a training sample consisted of three timesteps, during which the networks were fed an adapter image (*t*_1_), followed by a gray scale image (*t*_2_, pixel values of 0.5) and a test image (*t*_3_), with input size of 1 × 28 × 28 (channels × height 235 × width) for all timesteps. Since the adapter and test image are here each presented for only a single timestep, this set-up assumes no accumulation of adaptation within the presentation of adapter itself, which we found to reduce the model’s ability to capture the behavioral performance of the human participants in our experiment. To address this, we also presented networks with longer input sequences, both during model training and testing, using a 3:1 adapter-to-test duration ratio which mirrored the temporal structure of the visual stimuli shown to human participants (1500:500 ms, adapter:test). Sequence length was varied such that the test image was presented for a minimum of two timesteps and included sequences of adapter (*A*), blank (*B*) and test (*T*) of either 9 (*AAAAAABTT*), 13 (*AAAAAAAAABTTT*), 17 (*AAAAAAAAAAAABTTTT*) and 21 (*AAAAAAAAAAAAAAABTTTTT*) images. For all sequences, pixel values were clamped to the range [0, 1] to reduce numerical complexity and increase computational efficiency.

### Neural network training

All networks were implemented using Pytorch (Paszke et al., 2019) with standard convolutions and linear layers. Activations from the last time step were used for the classification, computation of the loss and weight updates. The backbone of all DCNNs consisted of three convolutional layers, one fully connected and a readout layer (**Fig. 2D**, for the implementation see cnn feedforward.py). We define a linear response **L***_n_*(*t*) at time *t* as the output of a convolutional layer *n*:

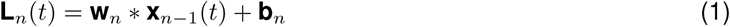

given the unit’s current input **x***_n_*_−1_(*t*), bottom-up convolutional weights **W***_n_* and biases **b***_n_*, whereby the convolution operation is represented as ∗. We trained five different model architectures. For the baseline model, no adaptation mechanism was added such that all images were processed independently over time. We then implemented and compared three different adaptation mechanisms (see below) in the model convolutional layers, such that responses on a given time step were modulated by responses on the previous time steps. Corresponding parameters were implemented per layer and were optimized simultaneously with regular DCNN parameters. Networks were trained using noise-embedded digits where the contrast was randomly varied between [0.1, 1] for 5 epochs (with 60000 images per epoch). To assess the robustness of our findings (Mehrer et al., 2020), 5 instances of DCNNs were trained from random initializations for each training length, resulting in a total of 20 trained models (4 model architectures x 5 initializations). Models were trained using the Adam optimizer, a learning rate of 0.001, the cross entropy using stochastic gradient descent with a batch size of 100. After the last convolution, 50% dropout was applied. Code used for model training can be found at model train.py. An overview of the number of parameters for each network architecture and dataset is provided in **Supplementary Table 1** and the temporal adaptation parameter values obtained after training are reported in **Supplementary Figure 1**.

### Temporal adaptation mechanisms

#### Additive suppression

We implemented an additive suppression mechanism that emerges from intrinsic properties of individual units (**Fig. 2E**, *left*), as introduced previously by Vinken et al. (2020). Here, each unit *i* in the network has an exponentially decaying adaptation state, *s_t_*, which is updated at each time step *t*, based on its previous state *s*(*t* − 1) and the previous response *r*(*t* − 1) as follows:

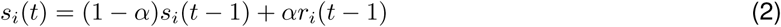

where *α* is a constant determining the time scale of the decay. This adaptation state is then subtracted from the linear response before applying the activation function *ϕ*.

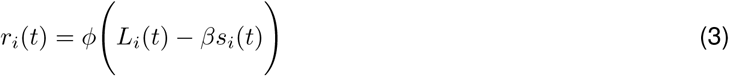

where *β* is a constant scaling the amount of suppression. For *β >* 0, these updating rules result in an exponentially decaying response for constant input that recovers in case of no input.

#### Recurrent interactions

In addition to temporal adaptation arising from intrinsic biophysical mechanisms, temporal adaptation phenomena have also been proposed to be the result of recurrent interactions. Here, we incorporated temporal adaptation via lateral recurrence using a method adapted from Vinken et al. (2020), inspired by computational models which implemented adaptation by changing recurrent interactions between orientation tuned channels (Felsen et al., 2002; Teich and Qian, 2003; Westrick et al., 2016):

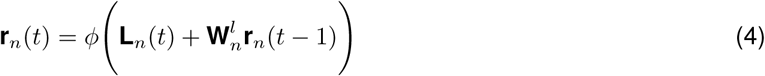

where 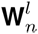 are the lateral weights (**Fig. 2E**, *right*) consisting of 32 kernels of size 1 × 1 × 32 (stride = 1).

#### Divisive normalization

Divisive normalization was originally proposed by Heeger (1992) and has been studied extensively in the spatial domain, where the response *r_i_* of neuron *i* is modelled as:

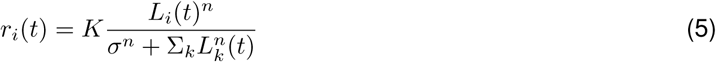

where *K* determines the maximum attainable response, *n* is an exponent, *σ* a semi-saturation constant and 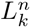 the sum of the activations of neighbouring neurons. Here, we transition to the temporal domain and implement a mathematical framework originally formulated in Heeger (1992, 1993), which defines a neuron’s response *r_i_* recursively over time (**Fig. 2F**). For each unit *i* in the network the response at timepoint *t* is updated before applying the rectifier activation function *ϕ* such that:

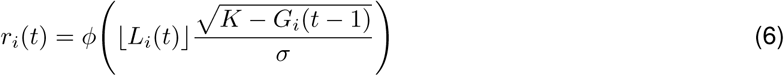

where *G*(*t* − 1) is a temporal feedback signal from the previous time step which is updated based on its previous state and the current response:

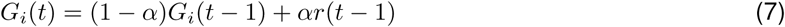

where *α* determines the time scale of the feedback signal. This multiplicative feedback signal results in divisive suppression (for details, see Heeger 1993, Appendix A).

### DCNN decoding analysis

To measure the amount of explicit information about the object present in DCNN activations and to compare with the results obtained from object decoding based on ERP responses, we also performed a decoding analysis on the DCNN unit activations of the last timestep (presentation of the test image) separately for each layer and contrast level. We applied a five-fold cross-validation, where each classifier was trained with 800 images (using the same type of classifier as was used for the decoding analysis of the ERPs, i.e. `sklearn`.`svm`.`SVC()`) and tested on 200 images (all randomly drawn from the test set). This procedure was repeated five times after which accuracies were averaged over DCNN instances.

## Data and code availability

The EEG data, behavioral responses, and stimuli are openly available on https://osf.io/ukqhg/. All code used for the purpose of this paper can be found at the GitHub repository https://github.com/ABra1993/tAdaptation_EEG.git.

## Acknowledgements

This work was supported by a MacGillavry Fellowship to IIAG.

## Supplementary information

**Supplementary Table 1:** Number of trainable parameters per model. Shown are the number of parameters for a feedforward model without (baseline) and with a temporal adaptation, arising from intrinsic (int) or recurrent (rec) mechanisms.

| model | # parameters |
| --- | --- |
| baseline | 178.058 |
| int. add. | 178.064 |
| lat. rec. | 181.226 |
| div. norm | 178.067 |

**Supplementary Figure 1:**
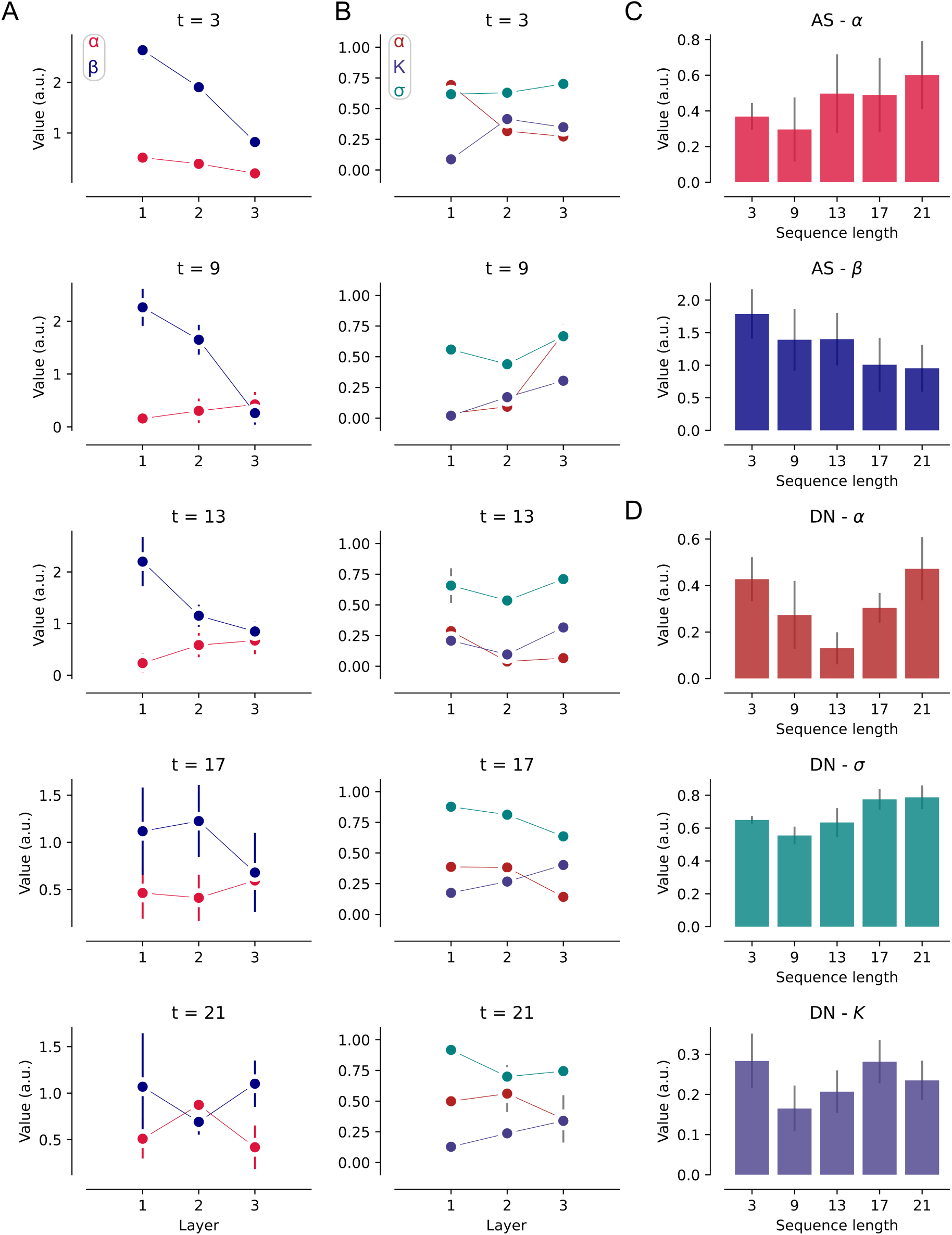
Temporal adaptation parameters for DCNNs with additive suppression and divisive normalization. **A**: Trained parameter values for each convolutional layer for DCNNs with additive suppression, including *α* and *β*. Each row depicts a different sequence length and error bars depict SEM across network initializations (*n* = 5). **B**: Same as panel A for DCNNs with divisive normalization, including the temporal adaptation parameters *α*, *K* and *σ*. **C**: Average parameter values for DCNNs with additive suppression averaged across the convolutional layers. Errors bars depict SEM across network initializations (*n* = 5). **D**: Same as panel C for DCNNs with divisive normalization. This figure can be reproduced by mkSFigure1.py.

**Supplementary Figure 2:**
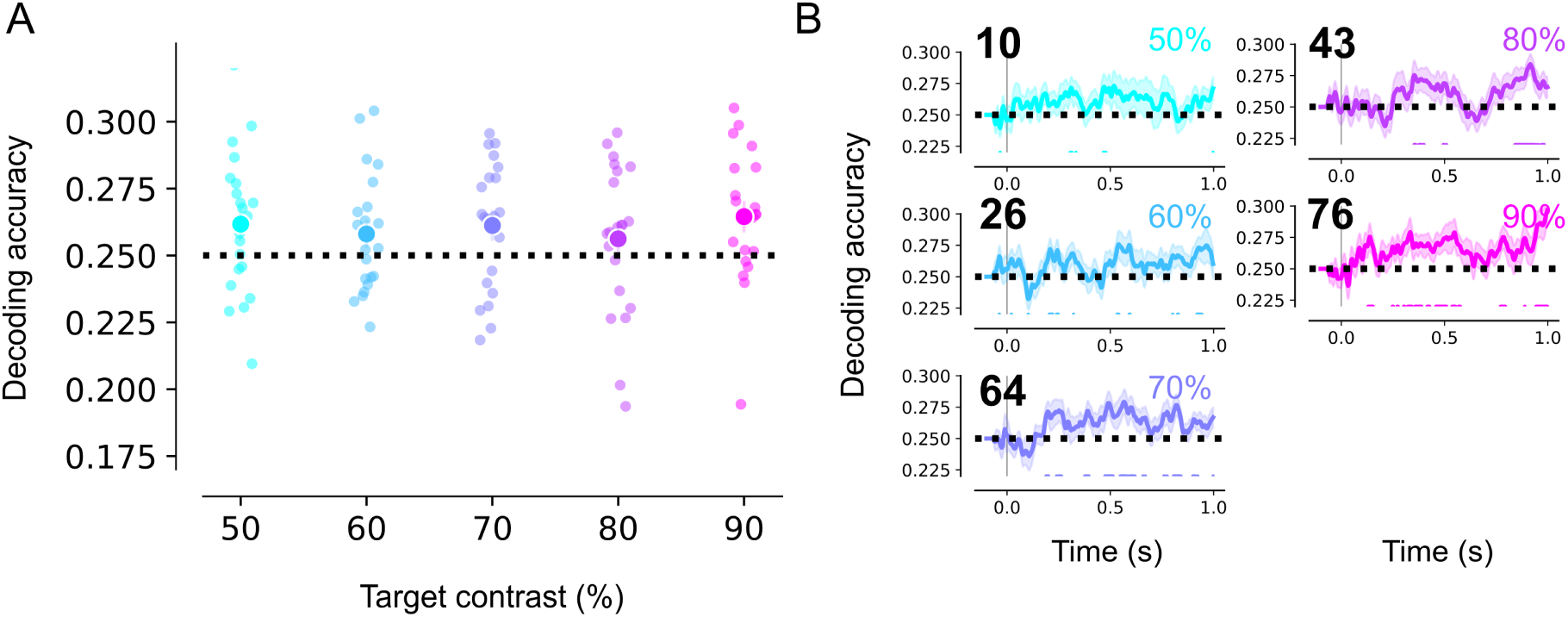
Object-related representations in the neural data per object contrast. **A**: Average decoding accuracy for predicting the presented object class based on evoked potentials for test images varying in object contrast level. Decoding accuracy’s are averaged for [0, 1]s time window after stimulus onset. **B**: Decoding accuracy for the different object contrast levels, with the number of time points for which decoding accuracy was significantly different from chance level (i.e. 25%) noted on the top left. Shaded regions depict the SEM across subjects. This figure can be reproduced by mkFigure7_SFig2.ipynb.

**Supplementary Figure 3:**
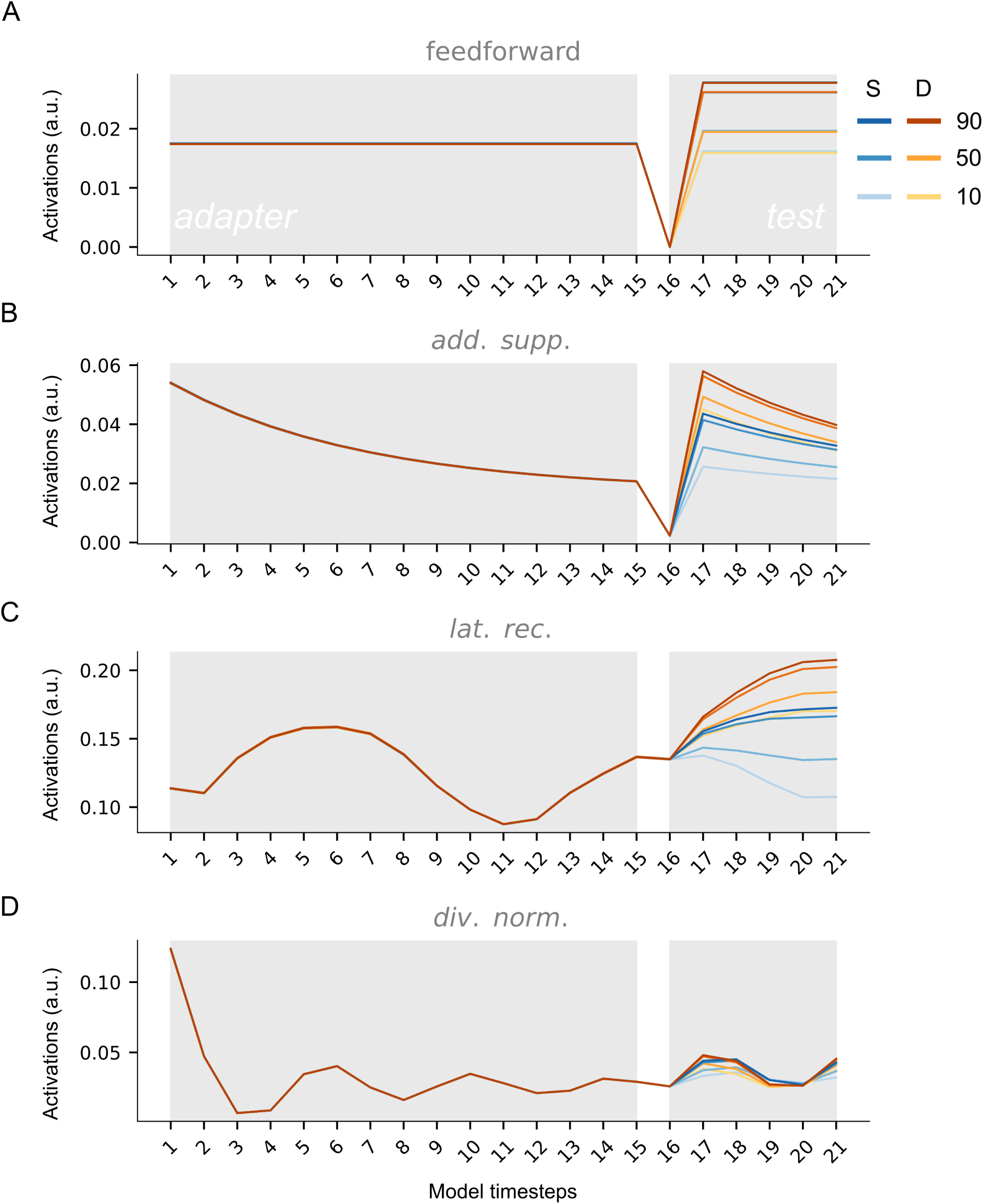
Model activations. **A**: Activations for the first convolutional layer for the test set for same (blue) and different (yellow) noise adapters for the first DCNN initialization without a temporal adaptation mechanism. Results are shown for three different object contrast levels (%), including 10, 50, 90. **B-D**: Same as panel A for a DCNN initialization endowed with a temporal adaptation mechanism, including additive suppression (B), lateral recurrence (C) and divisive normalization (D). This figure can be reproduced by mkSFigure3-7.py.

**Supplementary Figure 4:**
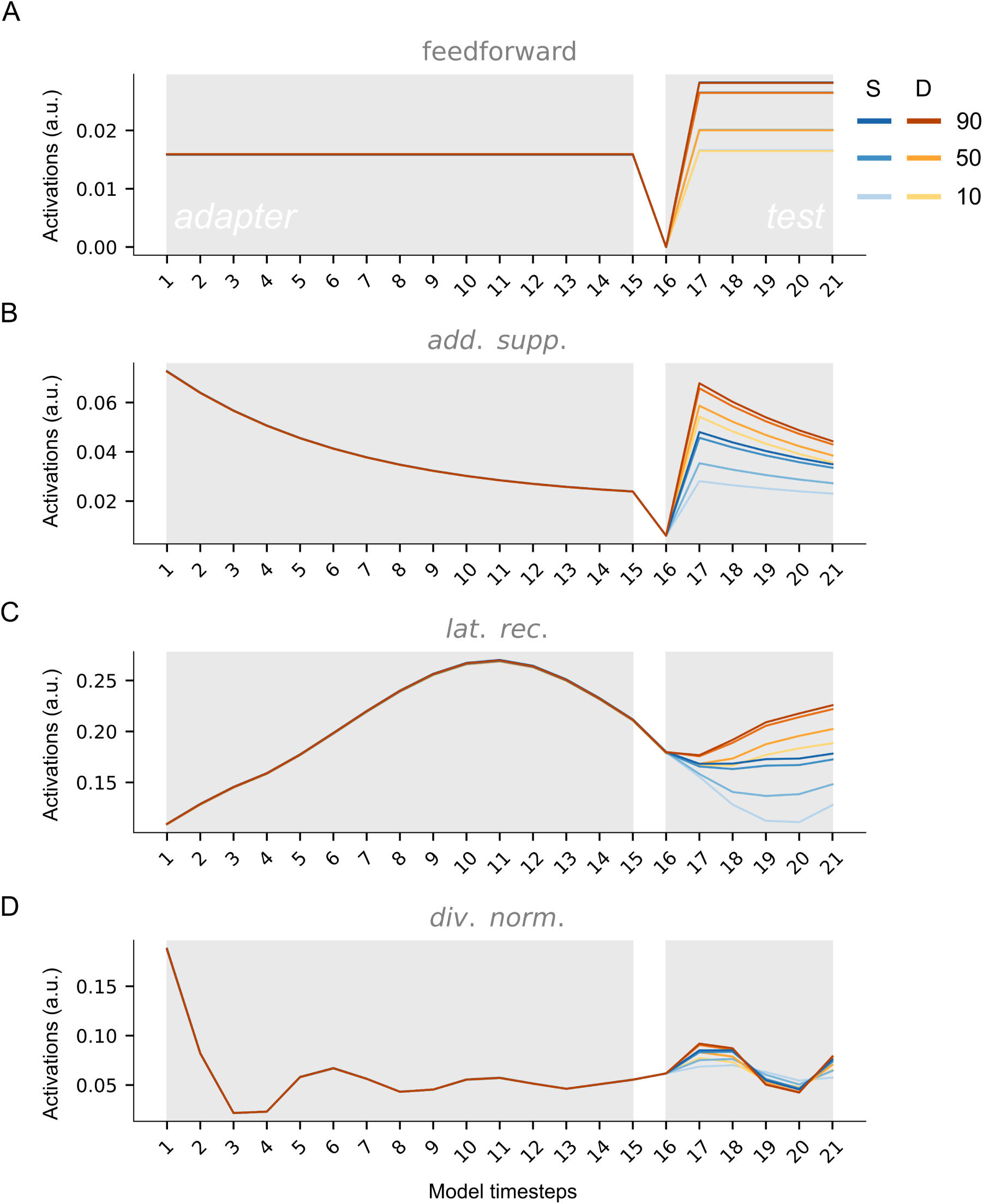
Model activations. **A**: Activations for the first convolutional layer for the test set for same (blue) and different (yellow) noise adapters for the second DCNN initialization without a temporal adaptation mechanism. Results are shown for three different object contrast levels (%), including 10, 50, 90. **B-D**: Same as panel A for a DCNN initialization endowed with a temporal adaptation mechanism, including additive suppression (B), lateral recurrence (C) and divisive normalization (D). This figure can be reproduced by mkSFigure3-7.py.

**Supplementary Figure 5:**
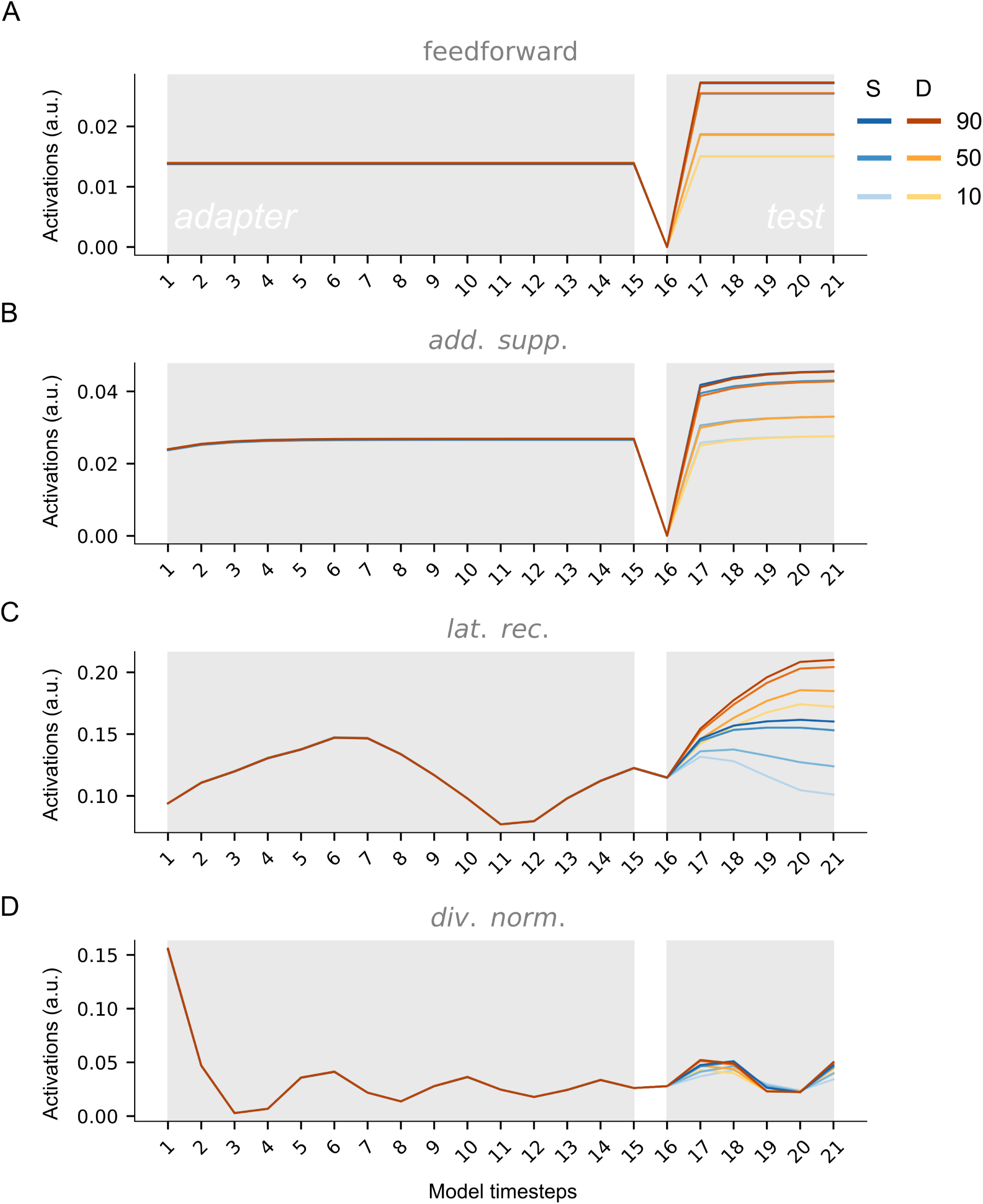
Model activations. **A**: Activations for the first convolutional layer for the test set for same (blue) and different (yellow) noise adapters for the third DCNN initialization without a temporal adaptation mechanism. Results are shown for three different object contrast levels (%), including 10, 50, 90. **B-D**: Same as panel A for a DCNN initialization endowed with a temporal adaptation mechanism, including additive suppression (B), lateral recurrence (C) and divisive normalization (D). This figure can be reproduced by mkSFigure3-7.py.

**Supplementary Figure 6:**
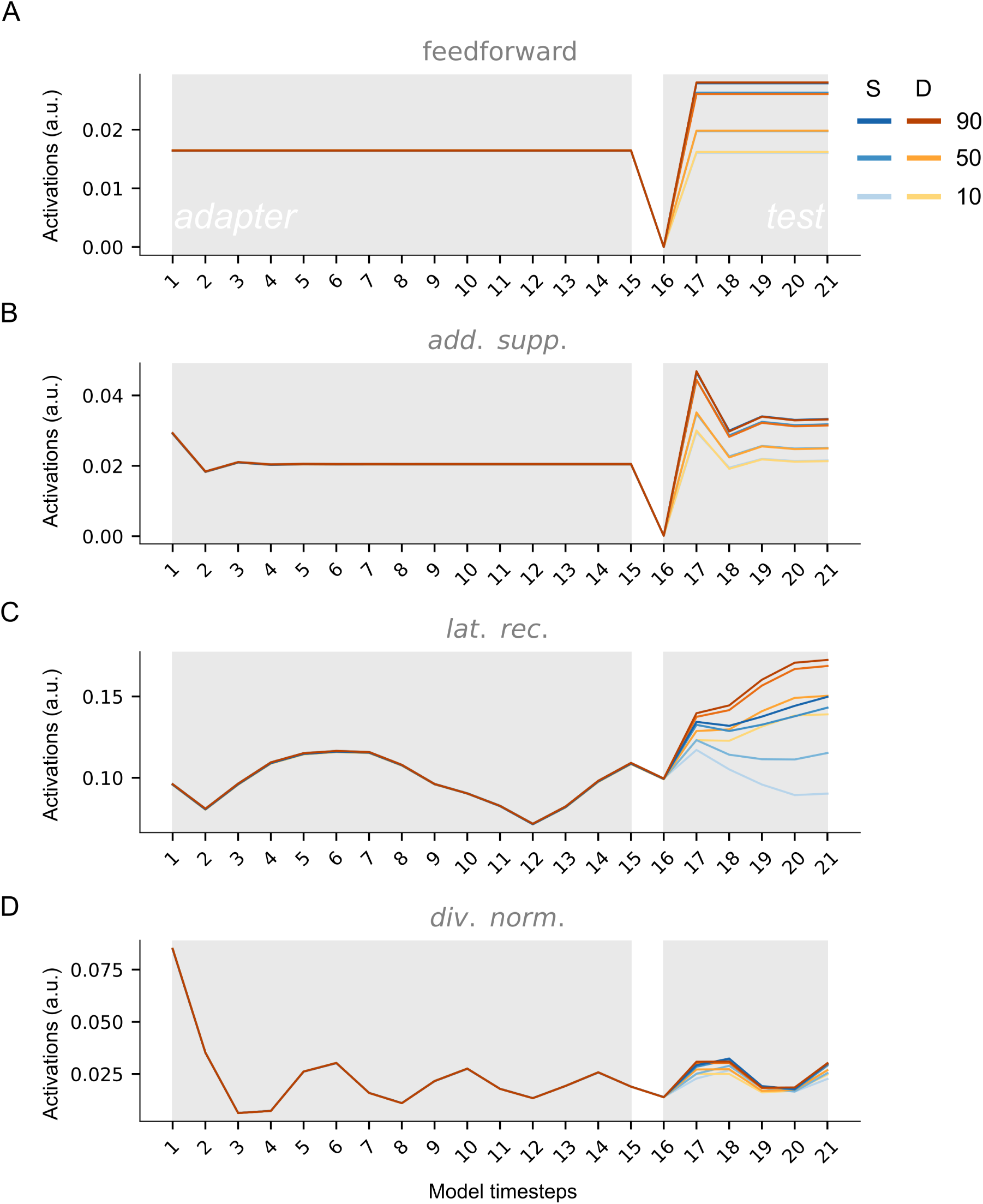
Model activations. **A**: Activations for the first convolutional layer for the test set for same (blue) and different (yellow) noise adapters for the fourth DCNN initialization without a temporal adaptation mechanism. Results are shown for three different object contrast levels (%), including 10, 50, 90. **B-D**: Same as panel A for a DCNN initialization endowed with a temporal adaptation mechanism, including additive suppression (B), lateral recurrence (C) and divisive normalization (D). This figure can be reproduced by mkSFigure3-7.py.

**Supplementary Figure 7:**
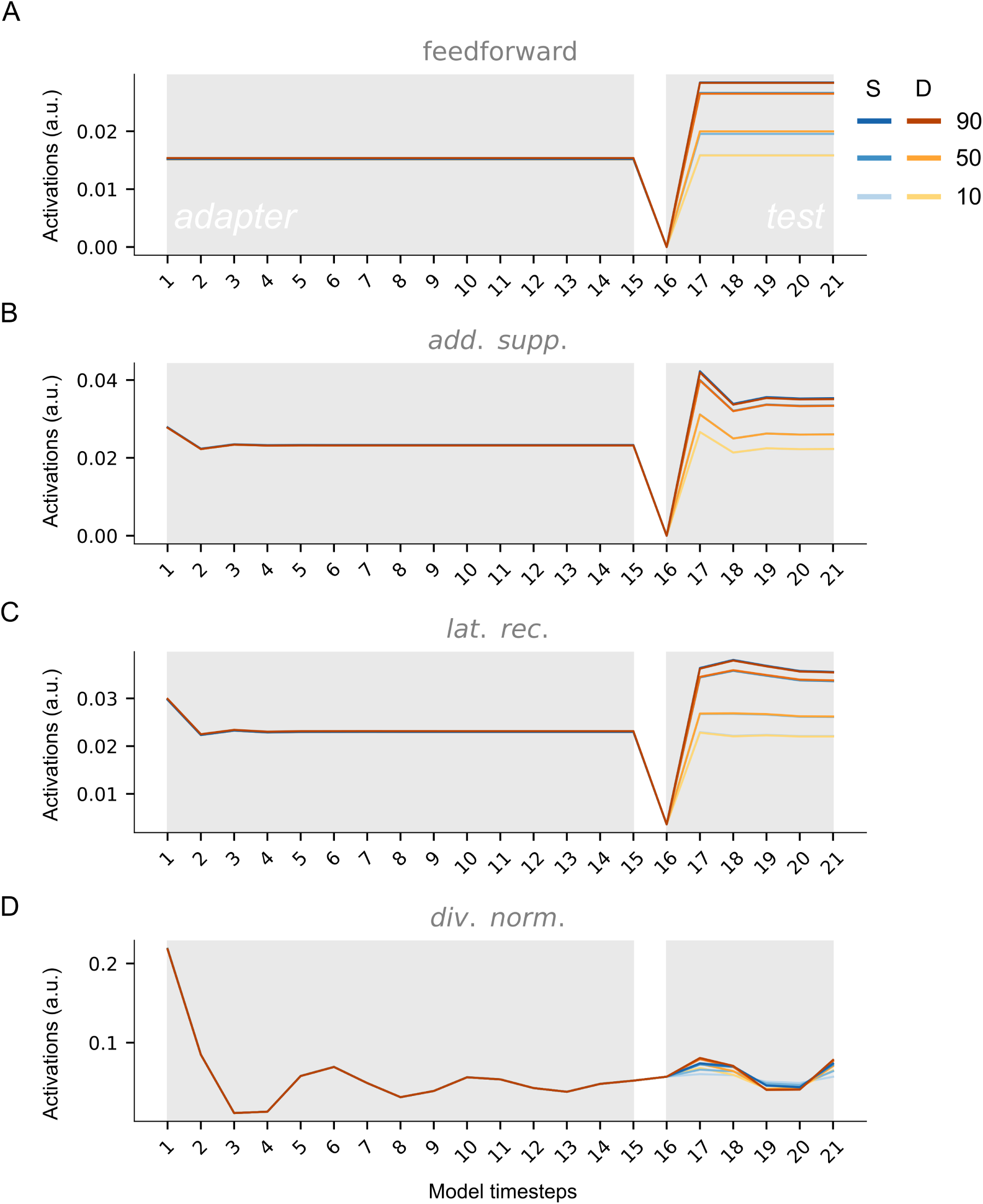
Model activations. **A**: Activations for the first convolutional layer for the test set for same (blue) and different (yellow) noise adapters for the fifth DCNN initialization without a temporal adaptation mechanism. Results are shown for three different object contrast levels (%), including 10, 50, 90. **B-D**: Same as panel A for a DCNN initialization endowed with a temporal adaptation mechanism, including additive suppression (B), lateral recurrence (C) and divisive normalization (D). This figure can be reproduced by mkSFigure3-7.py.

**Supplementary Figure 8:**
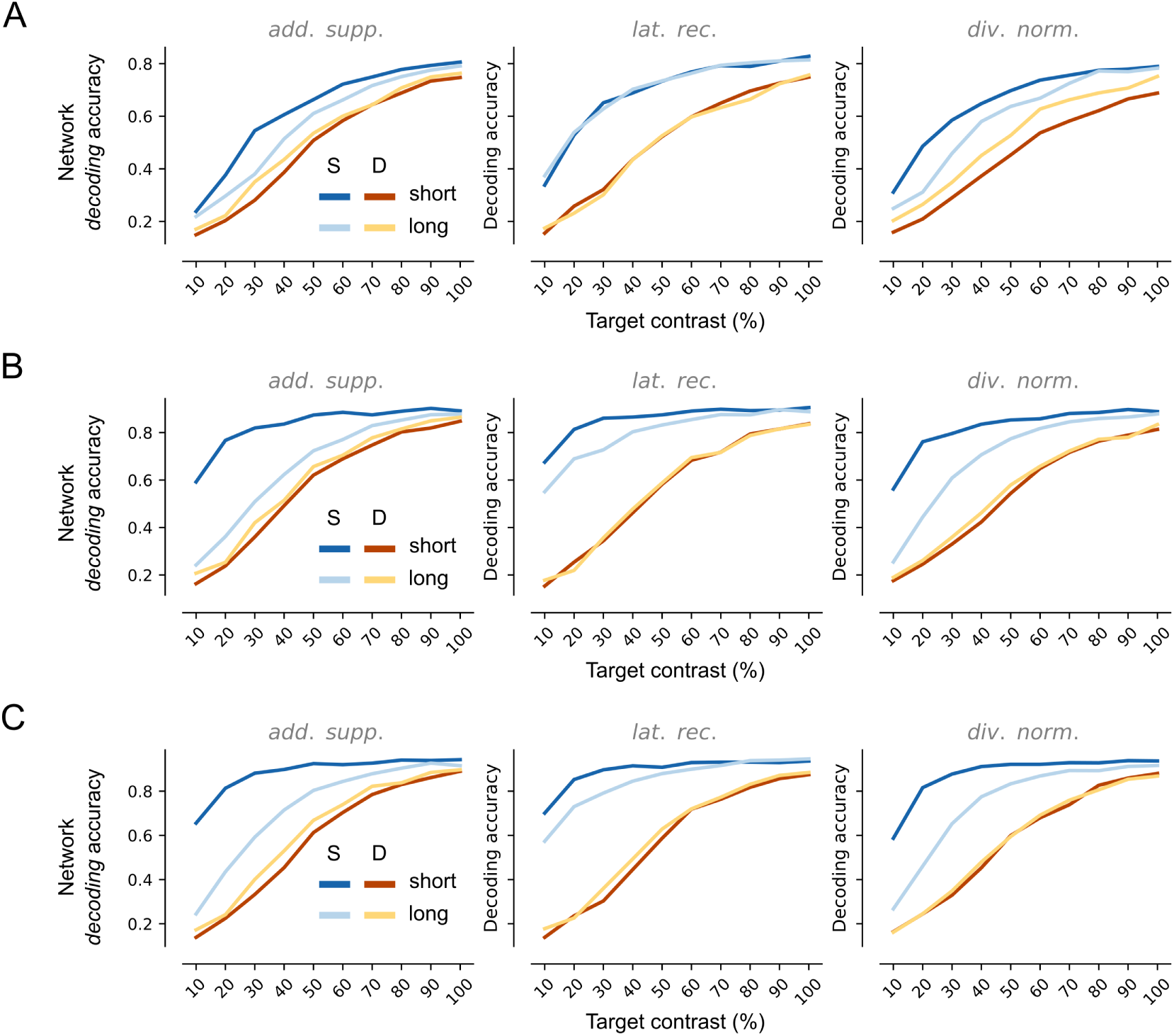
Decoding accuracy for the convolutional layers across adapter types. **A**: Decoding accuracy for the first convolutional layer for the test set for same (blue) and different (yellow) noise adapters for DCNNs endowed with one of three temporal adaptation mechanisms (from left to right): additive suppression, lateral recurrence and divisive normalization. Input sequences varied in length, including short (*ABT*) and long (*AAAAAAAAAAAAAAABTTTTT*) sequences. **B-C**: Same as panel A for the second (B) and third (C) convolutional layer.This figure can be reproduced by mkFigure8E_SFig8.py.

